# Non-canonical nuclear function of glutaminase cooperates with Wnt signaling to drive EMT during neural crest development

**DOI:** 10.1101/2024.04.17.589887

**Authors:** Nioosha Nekooie Marnany, Alwyn Dady, Frédéric Relaix, Roberto Motterlini, Roberta Foresti, Sylvie Dufour, Jean-Loup Duband

## Abstract

Metabolic reprograming has been linked to epithelial-to-mesenchymal transition (EMT) in cancer cells, but how it influences EMT in normal cells remains largely unknown. Here we explored how metabolism impacts delamination and migration of avian trunk neural crest cells, an important stem cell population of the vertebrate embryo. We report that delamination exhibits a quiescent metabolic phenotype whereas migration is characterized by OXPHOS-driven metabolism coupled to distinct expression of metabolic, EMT and developmental genes. While glucose and glutamine are required for delamination and migration, we uncover a novel role for glutamine and its catabolizing enzyme glutaminase in delamination. Specifically, glutamine is required for nuclear translocation of glutaminase, which interacts and cooperates with Wnt signaling to regulate EMT gene expression and cell cycle during delamination. Our data indicate that similarly to cancer cells, embryonic cells engage metabolic enzymes for non-canonical signaling functions to connect metabolism with EMT.

## INTRODUCTION

Epithelium-to-mesenchyme transition (EMT) refers to the cellular program by which, through diverse scenarios, epithelial cells lose their compact, ordered organization to become individual, migrating cells. This process plays crucial roles during embryogenesis, morphogenesis, and tissue repair, but it can also adversely cause organ fibrosis and promote cancer progression and metastasis [1]. EMT is context dependent and triggered by numerous inducers, whose signaling cascades all converge to the Snail, Zeb and Twist families of transcription factors (EMT-TFs) which control expression of genes for adhesion, polarity and cytoskeleton molecules [2].

Because it establishes a direct link between the environment and gene networks, cellular metabolism plays a key role in the unfolding of cell fate and function during EMT. Numerous studies uncovered the mutual relationships between EMT and metabolism, and the molecular mechanisms implicated are being unraveled in cancer cells [3-6]. Metabolic reprograming occurs often as a consequence of EMT during tumorigenesis, enabling cells to adapt to environmental changes. Thus, EMT inducers can alter the cellular metabolic activity [7] and EMT-TFs can modify expression of metabolic genes, profoundly affecting the glycolytic flux, oxygen consumption, or rewiring glycolysis to the pentose phosphate (PP) pathway [8, 9]. Conversely, changes in the metabolic activity of cancer cells favor EMT. A well-described example of this phenomenon is the hypoxia found in carcinoma that promotes anaerobic glycolysis and activates the hypoxic-inducible factor 1, a potent regulator of EMT-TFs [10, 11]. Another well-known metabolic process causing EMT in tumor cells is the Warburg effect in which energy derives mainly from aerobic glycolysis and less from mitochondrial oxidative phosphorylation (OXPHOS) [12, 13]. The mechanisms by which metabolic activity induces EMT in cancer cells are extremely diverse. They may rely on the accumulation of metabolites generated in mitochondria through the tricarboxylic acids (TCA) cycle, which serve as cofactors or substrates of histone modifiers involved in epigenetic regulations [14]. Alternatively, EMT can result from non-canonical functions of glycolytic enzymes, either through their release into the microenvironment where they act as autocrine factors inducing EMT-TFs or via their translocation into the nucleus where they exert a transcriptional regulatory activity [15-17]. However, the diverse molecular processes capable of inducing EMT along with the intrinsic heterogeneity of tumor cells make it difficult to decipher which metabolic events are decisive in EMT [6]. Moreover, most metabolic alterations found in cancer cells result from extensive genomic alterations and mitochondrial dysfunctions. Hence the need to investigate those mechanisms in normal cells that undergo EMT in a predictable stereotyped fashion.

Neural crest cells (NCCs) of the vertebrate embryo represent a powerful model system to tackle this question [18, 19]. Delamination, an EMT-related process, constitutes the founding event of this stem cell population originating from the neural tube (NT), allowing the release of nascent NCCs from their niche into the surroundings where they undergo migration and disperse away until differentiation [20-22]. Owing to this process, NCCs give rise to a wide array of cell types throughout the body, from cephalic skeletal tissues to melanocytes and peripheral neurons and glia. Recent studies provided evidence that glucose metabolism plays an important role in NCC delamination and migration. In a similar fashion to cancer cells, NCCs at cranial levels rely primarily on aerobic glycolysis to undergo EMT with Yap/Tead signaling serving as the intermediate between glycolysis and activation of EMT-TFs [23]. Unlike their cranial counterparts, trunk NCCs display mitochondrial-dependent glucose oxidation and mobilize several metabolic pathways downstream glucose uptake to execute their developmental program [24]. Interestingly, trunk and cranial NCC delamination events are intrinsically different in their kinetics, cellular processes, and in the molecular players recruited [20, 25, 26], thus raising the intriguing question that metabolic activity might determine the scenario by which cells undergo EMT.

Here, using *in vivo* and *in vitro* strategies, we explored how metabolism drives transition from delamination to migration in trunk NCCs. We report that delamination and migration differ in their metabolic requirements in relation with the developmental gene networks involved. We show that beside glucose, glutamine is required for NCC EMT and that glutaminase (GLS), the enzyme that catalyzes glutamine transformation into glutamate, plays a major role in this process. Strikingly, GLS function in delamination is independent of its enzymatic activity and is instead related to its nuclear translocation, suggesting a non-canonical gene regulatory function. In fact we find that GLS cooperates with β-catenin, a major player of Wnt signaling, to control expression of EMT-TFs and cell cycle during NCC delamination. Thus, our data indicate that embryonic cells share similar molecular processes with cancer cells, connecting metabolic events with gene regulation during EMT.

## RESULTS

To examine the metabolic processes involved in trunk NCC delamination at the transition between their pre-migratory and migratory states, we used the avian embryo, a model in which the spatiotemporal features of delamination have been precisely mapped and characterized [27, 28]. Delaminating and migrating NCCs can be discriminated using specific markers. Expressions of the EMT-TFs *Snail-2* and *Foxd-3* peak at onset of EMT in synchrony with downregulation of the cell adhesion molecule *Cadherin-6B*; these markers gradually decline at initiation of migration, while expression of *Sox-10*, a marker of migration, increases (Fig. 1A; Extended data Fig.1A) [29-32]. In the thoracic region, delamination of the NCC population occurs over a 15-h period from stage 12 of Hamburger and Hamilton (HH12; [33]) up to stage 15. At HH12, pre-migratory NCCs express EMT markers and are about to undergo delamination. At HH13, delamination culminates and, at stage HH14-15, while delamination is still underway, the already-delaminated NCCs start migrating (Fig. 1A; Extended data Fig.1A). NCC delamination can be investigated experimentally *in vivo* [34] and can be mimicked *in vitro* under defined cell culture conditions [35]. Therefore, this system allows to analyze the process of delamination and its switch to migration and to decipher the individual and collective behaviors of NCCs, thereby providing a comprehensive spatiotemporal survey of the underlying metabolic events.

**Figure 1:**
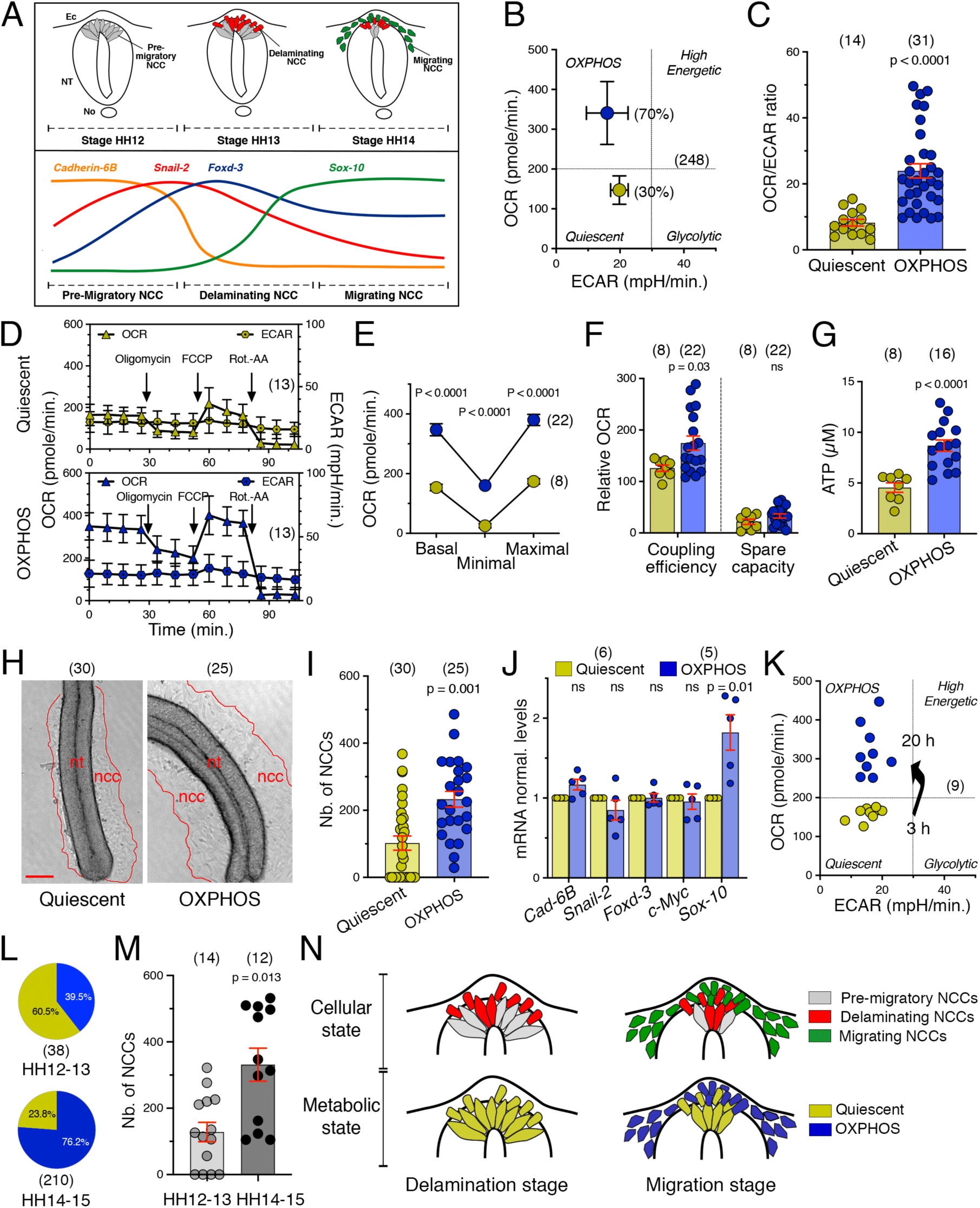
NCC transition from delamination to migration is coupled to metabolic switch. (A) Schematic representation of the gene regulatory network controlling NCC delamination and migration in the trunk region of the avian embryo. The top diagram depicts transverse views through the NT at the thoracic level at stages HH12, HH13 and HH14, showing the pre-migratory (grey), delaminating (red), and migrating (green) NCC populations. ec, ectoderm; no, notochord; nt, neural tube. The bottom diagram depicts the expression profiles of markers of NCC delamination and migration over time. (B) Energetic maps of the two metabolic profiles, quiescent (yellow) and OXPHOS (blue), obtained from the analysis of a large panel of explants at HH12-13 to HH14-15 cultured for 3 h in medium with glucose and glutamine. Data represent the mean values ± s.e.m. of OCR and ECAR. (C) Scatter dot plot of the OCR/ECAR ratio (mean ± s.e.m.) in NT explants with quiescent and OXPHOS profiles. (D) Mitochondrial stress assays of individual trunk NT explants with quiescent (top) and OXPHOS (bottom) profiles showing OCR and ECAR changes over time. Data were collected from at least three independent experiments. (E-G) Metabolic parameters (mean values ± s.e.m.) of NT explants with quiescent and OXPHOS profiles of: basal, minimal, and maximal OCR (E); coupling efficiency and spare respiratory capacity (F); and ATP levels (G). (H) Phase contrast images of NT explants with quiescent and OXPHOS profiles after 3 h in culture. The outermost boundary of the NCC outgrowth on either side of the NT is delineated in red. ncc, neural crest cells; nt, neural tube. Bar = 100 µm. (I) Scatter dot plots of the number of migrating NCCs per explants (mean ± s.e.m.) with quiescent and OXPHOS profiles after 3 h of culture. (J) Scatter dot plot of normalized mRNAs levels (mean ± s.e.m.) measured by qRT-PCR of delamination and migration markers in NT explants with quiescent and OXPHOS profiles. *c-Myc*, a gene expressed by delaminating and migrating NCCs but not implicated in EMT [52], was used as an internal control. (K) Energetic maps of 9 individual explants with an initial quiescent metabolic profile at 3 h of culture showing a switch to an OXPHOS profile after 20 h. (L) Proportions of quiescent and OXPHOS profiles in trunk NT explants at the thoracic level from HH12-13 and HH14-15 embryos, respectively. (M) Scatter dot plot of the number of migrating NCCs per explant (mean ± s.e.m.) at stages HH12-13 and HH14-15 after 3 h in culture. (N) Diagram depicting the coupling between the transition from delamination to migration and the metabolic shift from quiescence to OXPHOS in trunk NCCs. Images in H are from representative experiments. In panels B-I and K-M, (n) indicate the number of explants analyzed, and in J, (n) indicate the number of experiments with the measurements done in triplicates for each gene. In panels C, F, G, I and M, data were analyzed using unpaired two-tailed *t*-test. ns, not significantly different, P>0.05.

### NCC transition from delamination to migration is coupled to a metabolic switch

To study the bioenergetics profile of NT explants from embryos at stages encompassing the delamination process from HH12-13 to HH14-15, we measured at the onset of culture oxygen consumption rate (OCR) and extracellular acidification rate (ECAR) as readouts of mitochondrial respiration and glycolysis, using a Seahorse XF analyzer. For this aim, we optimized the methodological approach to individual NT explants for each measurement (Extended data Fig.1B), and we analyzed a large collection of samples (nearly 250) for greater accuracy and to circumvent the intrinsic variability in development progression among embryos.

Intriguingly, we consistently observed two subsets of explants with distinct metabolic states, a major one (70%) with an “OXPHOS-driven profile” characterized by high OCR, low ECAR and a high OCR/ECAR ratio, and a minor one (30%) with a “metabolically-quiescent profile” characterized by low OCR, low ECAR, and a low OCR/ECAR ratio (Fig. 1B,C). Using mitochondrial stress assays combined with measurements of intracellular ATP, we found that mitochondrial respiration was greater in explants with OXPHOS profile than in quiescent ones (Fig. 1D,E) and largely contributed to energy production (Fig. 1F,G). To determine whether these two metabolic profiles might be linked with NCC development, we assessed the cellular and molecular features of NCCs in the explants after metabolic profiling. Explants with an OXPHOS profile had a significantly greater capacity to produce large NCC outgrowths after 3 h of culture, exhibiting a higher number of migrating cells and stronger expression of the migration marker *Sox10* compared to the quiescent group (Fig. 1H-J). This suggests that explants with an OXPHOS profile may be at a more advanced development state compared to quiescent cells. We therefore performed a bioenergetics survey of individual explants over time in culture during the passage from delamination to migration, which occurs during the first day of culture. We found that explants exhibiting a quiescent profile at 3 h shifted to an OXPHOS profile after 20 h (Fig. 1K), while those with an OXPHOS profile at the beginning maintained the same metabolic activity later on, as reported previously [24]. This explains the metabolic heterogeneity observed in explants at 3 hours of culture. To further establish a correspondence between the stage of NCC development and their metabolic activity, we focused on embryos at stages covering either initiation (HH12-13) or termination (HH14-15) of delamination. The proportion of the metabolic profiles varied significantly between the developmental stages. Most explants at HH12-13 exhibited quiescent profiles while at HH14-15 a great majority of them displayed OXPHOS profiles (Fig. 1L). In addition, NTs from HH12-13 embryos cultured *in vitro* for 3 h produced fewer migrating NCCs than those from HH14-15 embryos (Fig. 1M).

Collectively, these data establish that the metabolic activity of NCCs is rewired from quiescence to active mitochondrial respiration at the transition phase between delamination and migration (Fig. 1N). Thus, metabolic profiling is instrumental to discriminate reliably NT explants at delamination and migration and investigate the molecular processes underlying the switch between the two stages.

### A change in glucose utilization accompanies NCC transition from delamination to migration

Metabolic switch is often related to changes in metabolite demand and utilization. To explore which metabolic programs are rewired during transition from delamination to migration, we focused first on glucose metabolism, as it is a main energy supplier. We previously demonstrated that glucose is required for trunk NCC development and that genes of several glycolytic enzymes are expressed in the embryo at the time of migration [24]. Here, after metabolic profiling of NT explants *in vitro*, we compared the mRNA levels at delamination and migration of the glucose transporter type-1 (*SLC2A1*/*GLUT-1*) and the enzymes phosphofructokinase-P (*PFK*), lactate dehydrogenase (*LDH*), and pyruvate dehydrogenase complex (*PDH*) (Fig. 2A). To demonstrate the relevance of our approach with the *in vivo* situation, data were compared with those obtained from extracts of the brachial region of embryos at HH12-13 and HH14 (see procedure Extended data Fig.1C). Consistent with a greater energy production, we found that except *LDH*, glycolytic enzymes were upregulated at migration compared with delamination (Fig. 2B).

**Figure 2:**
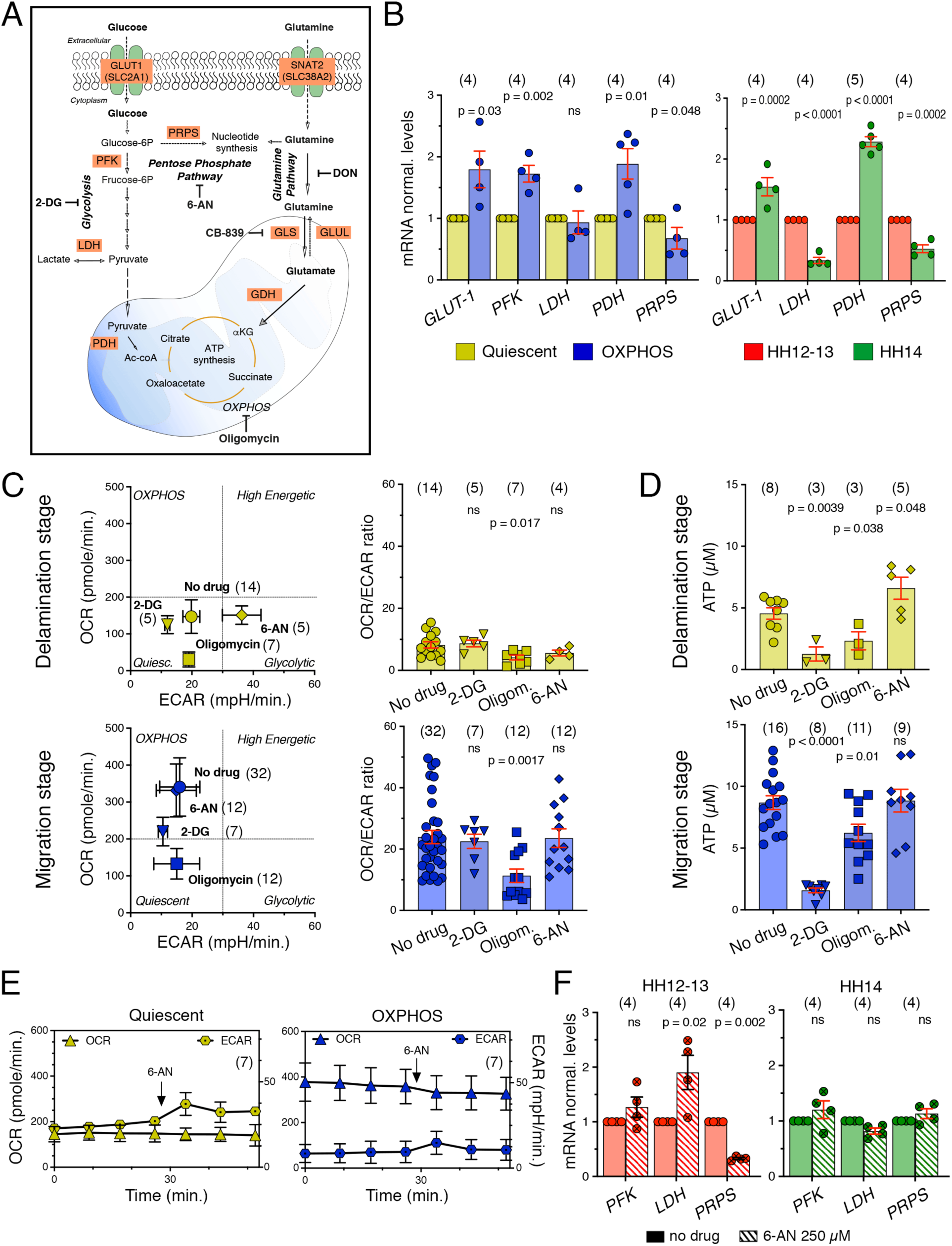
A change in glucose utilization accompanies NCC transition from delamination to migration. (A) Schematic representation of the most relevant metabolic pathways downstream of glucose and glutamine uptakes and their interconnections with the PP and OXPHOS pathways. The main enzymes within each pathway analyzed in this study as well as the specific metabolic inhibitors used are indicated. (B). Scatter dot plot of normalized mRNAs levels (mean ± s.e.m.) measured by qRT-PCR of genes encoding glycolytic enzymes in cultured NT explants with quiescent (yellow) and OXPHOS (blue) profiles (left) and in NTs collected from the trunk region of HH12-13 (red) and HH14 (green) embryos (right). (C, D) Effect of inhibitors of glycolysis (2-DG, triangles), OXPHOS (Oligomycin, squares), and PP pathways (6-AN, diamonds) compared with medium without drug (circles) on metabolic parameters (C) and ATP levels (D) of NT explants at the delamination (quiescent, top panels) and migration (OXPHOS, bottom panels) stages: energetic maps defined by OCR and ECAR as mean values ± s.e.m. (C, left panels); scatter dot plots of the OCR/ECAR ratio as mean ± s.e.m. (C, right panels). OCR and ECAR were recorded in individual explants before and after inhibitor treatment and ATP levels were measured after two additional hours in the presence of the inhibitor. (E) Changes in OCR and ECAR over time of individual trunk NT explants with quiescent (left) and OXPHOS (right) profiles before and after addition of 6-AN. Data were collected from at least three independent experiments. (F) Scatter dot plot of normalized mRNAs levels (mean ± s.e.m.) measured by qRT-PCR of *LDH, PFK* and *PRPS* in the trunk of embryos 5 h after 6-AN or vehicle (no-drug) injection. Embryos were injected either at HH12-13 (red) or at HH14 (green). In panel B left, (n) indicates the number of experiments with the measurements done in triplicates for each gene; in panels B right and F, (n) indicate the number of embryos, with the measurements done in triplicates for each gene. In C-E, (n) indicate the number of explants analyzed. Data in B and F were analyzed using unpaired two-tailed *t*-test. Data in C and D were analyzed using one-way ANOVA followed by Dunnett’s multiple comparison tests relative to the condition with quiescent profile. ns, not significantly different, P>0.05.

Next, we analyzed the effect of pharmacological inhibitors of glycolysis and OXPHOS, namely 2-deoxyglucose (2-DG) and oligomycin (Fig. 2A), on the metabolic activity of NT explants, by recording OCR and ECAR in individual explants and by measuring ATP levels. In explants at delamination, we observed that oligomycin decreased OCR and reduced ATP levels, while 2-DG did not affect OCR but decreased ATP (Fig. 2C,D). In contrast, consistent with our previous data [24], explants at migration exhibited a drastic decrease in OCR and ATP with 2-DG and oligomycin (Fig. 2C,D). This indicates that during delamination, rather than fueling glycolysis and OXPHOS, glucose might be utilized by alternative routes, such as the PP pathway, while during migration, NCCs use glucose oxidation and mitochondrial respiration as the main route for energy production. Consistent with this assumption, expression of the gene for the phosphoribosyl-pyrophosphate synthase (PRPS), a key enzyme of purines and pyrimidines synthesis in the PP pathway, was higher at delamination than migration phase (Fig. 2B). We therefore evaluated the effect of the PP pathway inhibitor 6-amminonicotinamide (6-AN) (Fig. 2A) on NCC metabolic activity. We found that in explants at delamination 6-AN had no impact on OCR but increased ECAR and ATP (Fig. 2C-E), suggesting that blocking the PP pathway re-diverts glucose towards glycolysis and lactate production. In contrast, in explants at the migration stage, 6-AN treatment changed neither the metabolic profile nor ATP levels (Fig. 2C-E). We next quantified the mRNA expression of enzymes of glucose metabolism in embryos at the time of NCC delamination and migration after *in ovo* injection of 6-AN (see procedure Extended data Fig.1D). We found that, at delamination 6-AN caused a significant reduction in *PRPS* expression and an increase in *LDH* with no effect on *PFK*, while no significant changes in these genes were detected during the migration phase (Fig. 2F). Altogether, these findings support a change in glucose utilization during NCC transition from delamination to migration.

### Glutamine metabolism is required for NCC delamination and migration

Because a metabolic switch may depend also on the type of metabolite used, we considered other metabolic pathways that may accompany NCC transition from delamination to migration. We focused on glutamine, a major contributor to macromolecules and ATP production known to cooperate with glucose in cancer cells [36]. First, we explored the expression patterns of key enzymes involved in glutamine metabolism (Fig. 2A) in the trunk of avian embryos, by in situ hybridization on whole mount embryos and on sections. mRNAs for *SNAT-2*, encoding one of the glutamine transporters in neural cells, glutaminase (*GLS*) and glutamate dehydrogenase (*GDH*) were all widely expressed at delamination and migration (Fig. 3A). At the tissue level, mRNAs were distributed throughout the NT and were also detectable in pre-migratory, delaminating and early-migrating NCCs (Fig. 3B). We also compared the expression levels of these enzymes in metabolically profiled explants cultured *in vitro* as well as in embryo extracts at HH12-13 and HH14. We found that *SNAT-2*, *GLS* and *GLUL* expressions remained constant throughout NCC progression from delamination to migration, while *GDH*, encoding the enzyme involved in glutamate conversion into α-ketoglutarate, was increased at migration (Fig. 3C). These data indicate that glutamine metabolism is active during both delamination and migration and suggest that, like for glucose, a change in glutamine utilization occurs during transition between the two stages.

**Figure 3:**
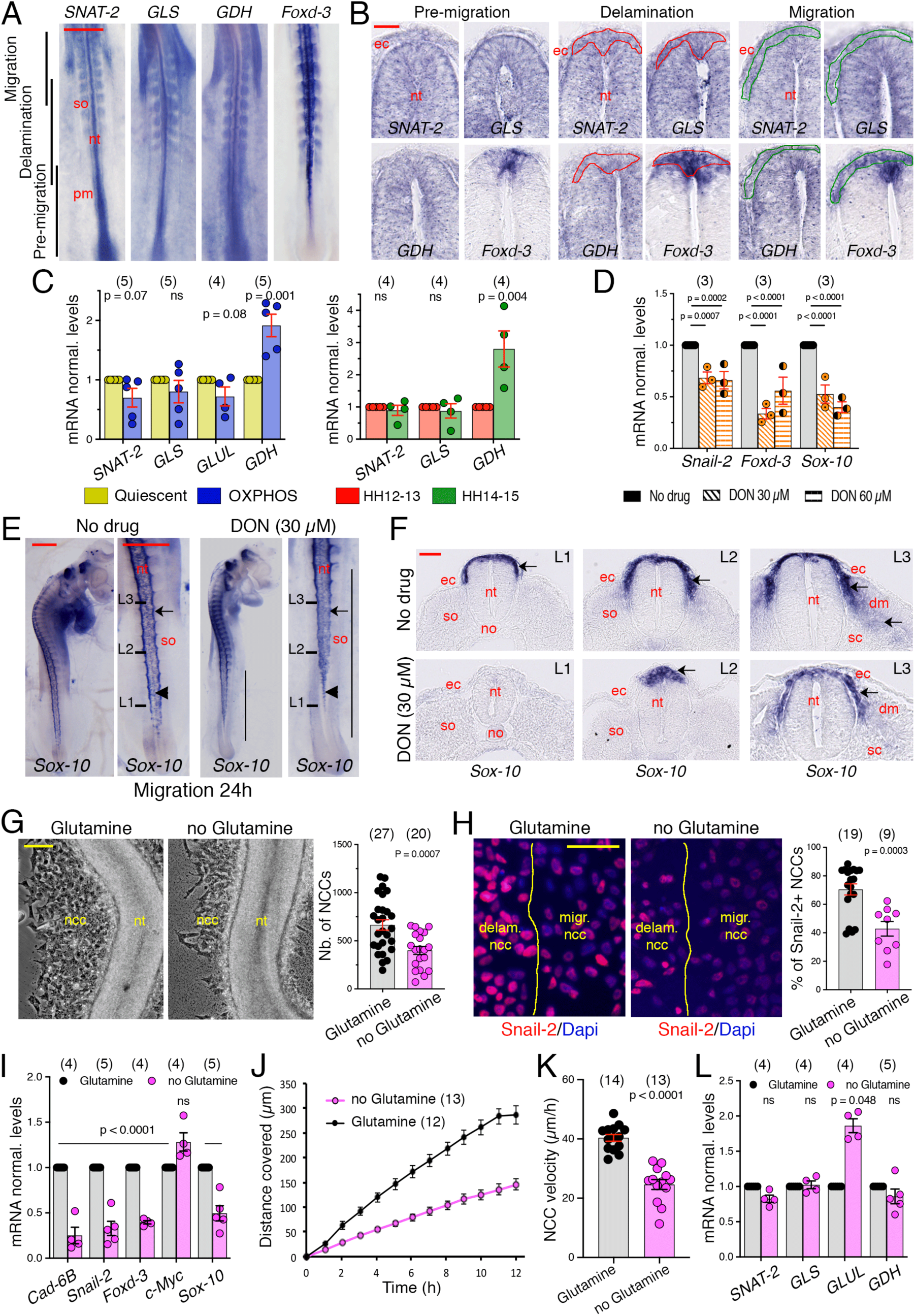
Glutamine metabolism is required for NCC delamination and migration. (A) Whole mount views of the trunk of HH13 quail embryos hybridized with probes for the players in glutamine metabolism *SNAT-2*, *GLS*, and *GDH*, and for *Foxd-3* used as a reference marker for the axial levels of NCC delamination and migration; anterior to the top. In situ hybridizations for each probe were done in triplicate. nt, neural tube; pm; unsegmented paraxial mesoderm; so, somite. Bar = 500 µm. (B) Cross-sections through the caudal half of whole-mount embryos hybridized with probes for *SNAT-2*, *GLS*, *GDH* and *Foxd-3* showing the dorsal NT at the levels of pre-migratory (left panels), delaminating (delineated in red; middle panels) and early-migrating (delineated in green; right panels) NCCs. ec, ectoderm, nt, neural tube. (C) Scatter dot plot of normalized mRNAs levels (mean ± s.e.m.) measured by qRT-PCR of *SNAT-2*, *GLS*, *GDH*, and *GLUL*, in cultured NT explants with quiescent (yellow) and OXPHOS (blue) profiles (left) and in NTs collected from the trunk region of HH12-13 (red) and HH14 (green) embryos (right). (D) Scatter dot plot of normalized mRNAs levels (mean ± s.e.m.) measured by qRT-PCR of *Snail-2, Foxd-3* and *Sox-10* in the trunk of embryos 5 h after DON or vehicle (no-drug) injection. (E) Whole mount views of embryos 24 h after DON or vehicle injection at HH12 hybridized with probes for *Sox-10*. For each condition, an overall view of the embryo is shown on the left (red bar = 250 µm) and a detailed view of the trunk region is shown on the right (red bar = 500 µm). Vertical bars delineate the axial levels where NCCs are defective. Arrowheads and arrows point at delaminating and migrating NCCs, respectively. L1, L2 and L3-labeled bars indicate the levels of the cross-sections shown in F. n = 10 for vehicle injection with 100% of non-affected embryos and n = 21 for DON injection with 90% of the embryos showing apparent defects in NCC delamination and migration. nt, neural tube; so, somite. (F) Cross-sections of whole-mount ISH for Sox-10 at distinct levels (indicated in E) through the trunk of embryos 24 h after DON or vehicle injection. Arrows point at NCCs. dm, dermamyotome; ec, ectoderm; no, notochord; nt, neural tube; sc, sclerotome; so, somite. Bar = 50 µm. (G) Left: Phase contrast images of HH12/13-14 trunk NT explants after 5 h in culture in medium containing both glucose and glutamine or without glutamine. ncc, neural crest cells; nt, neural tube. Bar = 100 µm. Right: Scatter dot plot with mean ± s.e.m. of the total number per explant of NCCs (both delaminating and migrating, visualized after removing of the NT explant). (H) Left: Immunofluorescence staining for Snail-2 (red) with Dapi visualization of nuclei (blue) in HH12/13-14 trunk NT explants after 5 h in culture in medium containing both glucose and glutamine or without glutamine. The NT was removed before immunostaining to visualize delaminating NCCs. The boundary between delaminating and migrating NCCs is delineated with a yellow line. Bar = 100 µm. Right: Scatter dot plot of the percentage of Snail-2-positive NCCs (both delaminating and migrating) per explant. (I) Scatter dot plot of normalized mRNAs levels (mean ± s.e.m) measured by qRT-PCR of NCC delamination and migration markers in HH12/13-14 explants cultured in medium with or without glutamine (black and magenta symbols, respectively). (J, K) Incidence of the lack of glutamine in the culture medium on NCC locomotion properties, as measured by video-microscopy using Metamorph software: Graph of the distance covered by the NCC population over time (J) and scatter dot plots of the NCC velocity (mean ± s.e.m.) throughout the duration of the experiment (K) in medium containing or not glutamine. Time 0 corresponds to the onset of recording 2-4 h after initiation of culture. Each dot in K corresponds to the mean velocity measured for 20 NCCs taken at the periphery of the outgrowth of a NT explant. (L) Scatter dot plot of normalized mRNAs levels (mean ± s.e.m) measured by qRT-PCR of enzymes of the glutamine pathway in NT explants cultured in medium with or without glutamine. Images in panels A, B, E, F, G and H are from representative experiments. In C left, I and L, (n) indicate the number of experiments, with measurements done in triplicate for each gene. In C right and D, (n) indicate the number of embryos, with measurements done in triplicate for each gene. In G, H, J and K, (n) indicate the number of explants analyzed. In C, I and L, data were analyzed using unpaired two-tailed *t*-test. In D, data were analyzed by two-way ANOVA relative to the condition of “No drug”. In G, H, J and K data were analyzed using one-way ANOVA followed by Dunnett’s multiple comparison tests relative to the condition in medium with glutamine. ns, not significantly different, P>0.05.

To assess the implication of glutamine metabolism in delamination and migration, we used the *in vivo* loss-of-function strategy (Extended data Fig.1D). DON, a glutamine analog that blocks all metabolic routes downstream of glutamine uptake (Fig. 2A), was injected in the caudal region of HH12 embryos. Analyses by qRT-PCR of the embryos 5 h following injection revealed a robust repression of *Foxd-3* and *Sox-10* and a more moderate decrease of *Snail-2* (Fig. 3D). Accordingly, DON strongly decreased NCC delamination and caused migration defects after 24 h, as assessed by the alteration of *Sox-10* pattern in the lower trunk (Fig. 3E). Spatiotemporal analyses of NCC distribution showed that cells were missing at the top of the NT in the caudal trunk and were less numerous and trapped dorsally instead of migrating ventrally in the mid-trunk (Fig. 3F). Therefore, consistent with the expression of glutamine enzymes throughout NCC delamination and migration, these results establish that glutamine is required for both events to occur.

In order to corroborate the role of glutamine, we analyzed the cellular and molecular responses of NT explants confronted *in vitro* with medium depleted of glutamine. We found that, similarly to DON effect, glutamine deprivation strongly impacted the numbers of NCCs (Fig. 3G) and of Snail-2-positive cells (Fig. 3H) and the expression of EMT-TFs (Fig. 3I). The absence of glutamine also reduced the expression of *Sox-10* (Fig. 3I), the outward expansion of the NCC population (Fig. 3J) and individual cell velocity (Fig. 3K). Interestingly, *GLUL*, encoding glutamine synthetase that converts glutamate into glutamine, was upregulated possibly due to a feedback adaptation (Fig. 3L). We also found by metabolic profiling that none of the NT explants displayed a quiescent profile typical of the delamination stage. Rather, despite a reduced capacity to undergo delamination and migration, the explants possessed an OXPHOS profile, mitochondrial functions, and ATP levels (Extended data Fig.2A-E) similar to those observed in NTs at migration (compare with Fig. 1B,C,E-G). Thus, it appears that glucose oxidation and mitochondrial respiration cannot compensate for the lack of glutamine, despite their robust capacity to provide energy. This result indicates that glutamine likely contributes to delamination and migration independently of energy production, suggesting the intriguing possibility that metabolic quiescence during delamination is related to glutamine metabolism.

### Glutaminase exhibits a non-canonical function during NCC delamination

To investigate how glutamine metabolism drives NCC delamination and migration, we targeted GLS, the enzyme involved in glutamine-to-glutamate conversion. To this aim, we injected *in ovo* prior to delamination at HH12 either the CB-839 compound, to block GLS activity (Fig. 2A) [37], or siRNAs to *GLS* to repress its expression. Both strategies resulted in severe alterations of NCC delamination and migration. Specifically, CB-839 significantly decreased *Foxd-3* and *Sox-10* mRNA levels (Fig. 4A), and strongly reduced the expression domain of *Foxd-3* in the dorsal NT at 5 h during delamination and of *Sox-10* at 24 h during migration (Fig. 4B,C). In addition, when applied onto NT explants in culture, CB-839 strongly diminished the numbers of NCCs and of Snail-2-expressing cells (Fig. 4D). Silencing *GLS* by siRNAs caused a dose-dependent reduction of its expression with an almost complete deletion at the highest dose (Fig. 4E). As observed with CB-839, *GLS* extinction repressed *Snail-2*, *Foxd-3* and *Sox-10* expression (Fig. 4E), accompanied by a severe reduction of NCC delamination and migration (Fig. 4F,G).

**Figure 4:**
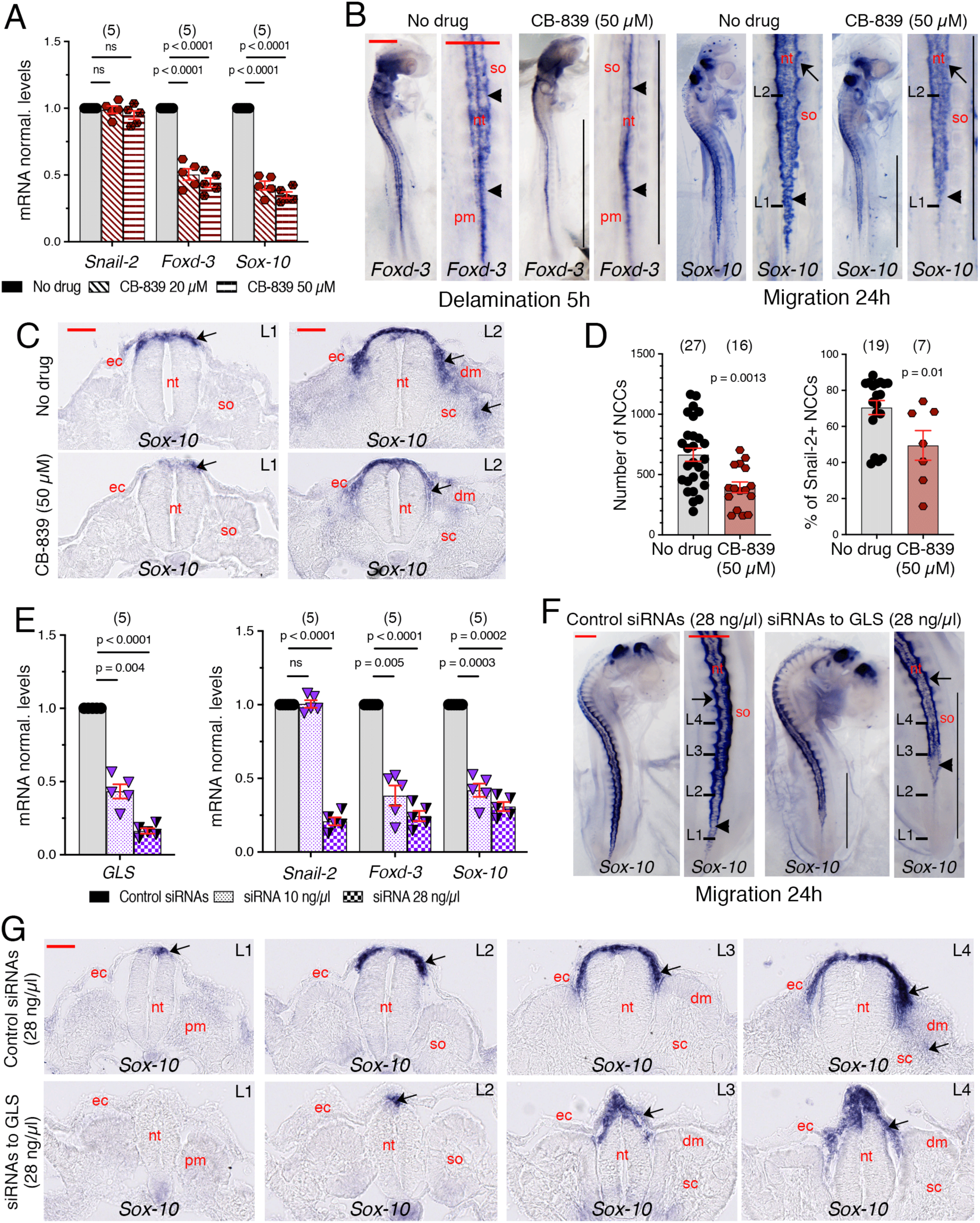
Glutaminase is required for NCC delamination and migration. (A) Scatter dot plot of normalized mRNAs levels (mean ± s.e.m) measured by qRT-PCR of markers of NCC delamination and migration in the trunk of embryos 5 h after CB-839 or vehicle (No-drug) injection. (B) Whole mount views of the trunk region of embryos 5 h (left panels) and 24 h (right panels) after CB-839 or vehicle injection, hybridized with probes for *Foxd-3* and *Sox-10*, respectively. For each condition, an overall view of the embryo is shown on the left (red bar = 250 µm) and a detailed view of the trunk region is shown on the right (red bar = 500 µm). Vertical bars delineate the axial levels where NCCs are defective. Arrowheads and arrows point at delaminating and migrating NCCs, respectively. L1 and L2-labeled bars indicate the levels of the cross-sections shown in C. n = 10 and 11 for vehicle injection with 100% normal embryos at 5 h and 24 h, respectively; and n = 7 and 11 for CB-839 injection with 71% and 73% of the embryos showing apparent defects in NCC delamination and migration at 5 h and 24 h, respectively. nt, neural tube; pm, unsegmented paraxial mesoderm; so, somite. (C) Cross-sections of whole-mount ISH for Sox-10 at different levels (indicated in B) through the trunk of embryos 24 h after CB-839 or vehicle injection. Arrows point at NCCs. dm, dermamyotome; ec, ectoderm; nt, neural tube; sc, sclerotome; so, somite. Bar = 50 µm. (D) Scatter dot plots of the total number of NCCs (left) and of the percentage of Snail-2-positive NCCs (right) per explant (mean ± s.e.m.) in the presence or not of CB-839 during 5 hours. (E) Scatter dot plot of normalized mRNAs levels (mean ± s.e.m) measured by qRT-PCR of *GLS* (left) and markers of NCC delamination and migration (right) in the trunk of embryos 5 h after injection of control siRNAs or of siRNAs to GLS. (F) Whole mount views of embryos 24 h after injection of control siRNAs or of siRNAs to GLS and hybridized with probes for *Sox-10*. For each condition, an overall view of the embryo is shown on the left (red bar = 250 µm) and a detailed view of the trunk region is shown on the right (red bar = 500 µm). Vertical bars delineate the axial levels where NCC development is defective. L1, L2, L3 and L4-labeled bars indicate the levels of the cross-sections shown in G. Arrowheads and arrows point at delaminating and migrating NCCs, respectively. n = 11 for injection of control siRNAs with 100% of non-affected embryos and n = 17 for injection of siRNAs to GLS with 88% of the embryos showing apparent defects in NCC delamination and migration. nt, neural tube; so, somite. (G) Cross-sections of whole-mount ISH for Sox-10 at distinct levels (indicated in F) through the trunk of embryos 24 h after injection of control siRNAs or of siRNAs against GLS. Arrows point at NCCs. dm, dermamyotome; ec, ectoderm; no, notochord; nt, neural tube; sc, sclerotome; so, somite. Bar = 50 µm. Images in panels B, C, F and G are from representative experiments. In A and E, (n) indicate the number of embryos with the measurements done in triplicates for each gene and in D, (n) indicate the number of explants analyzed. In A and E, data were analyzed by two-way ANOVA relative to the no drug condition. In D, data were analyzed using one-way ANOVA followed by Dunnett’s multiple comparison tests relative to the no drug condition. ns, not significantly different, P>0.05.

To further understand the role of GLS in these events, we compared the metabolic, cellular, and molecular responses of NT explants treated with CB-839 at delamination and migration phases. We first characterized the metabolic profiles of individual explants before and after addition of CB-839, and we quantified the expression of NCC markers and glutamine metabolism enzymes, along with ATP levels. Interestingly, GLS inhibition at delamination caused a significant decrease in *Snail-2*, *Foxd-3* and *Sox-10* expressions, but had no impact on OCR, ECAR and ATP levels (Fig. 5A,B,D). Conversely, at migration, CB-839 failed to alter the expression of NCC markers, but it decreased OCR and ATP production (Fig. 5A,B,D). Moreover, CB-839 differentially affected the expression of glutamine enzymes during delamination and migration (Fig. 5C). Notably, *GLUL* expression was reduced during delamination but not during migration. These data suggest that GLS may play distinct roles during delamination and migration.

**Figure 5:**
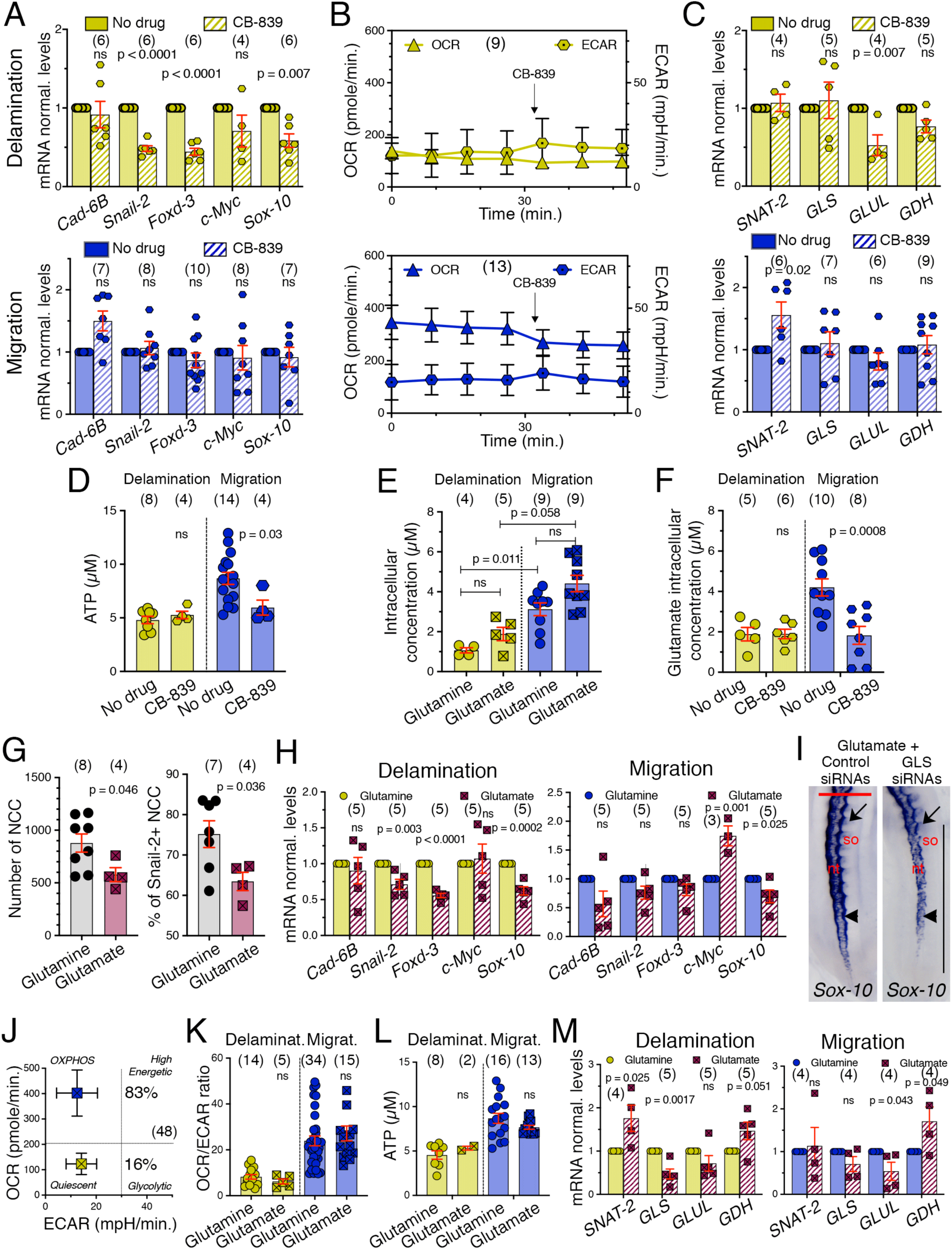
Glutaminase exhibits a non-canonical function during NCC delamination. (A) Effect of CB-839 (50 µM) on OCR and ECAR in individual trunk NT explants at the delamination (yellow; top) and migration (blue; bottom) stages. (B, C) Scatter dot plots of normalized mRNAs levels (mean ± s.e.m) measured by qRT-PCR of NCC delamination and migration markers (B) and players of the glutamine pathway (C) in trunk NT explants at the delamination (top) and migration (bottom) stages, in CB-839 (50 µM) or vehicle (No drug). (D) Scatter dot plot of ATP levels (mean ± s.e.m.) in explants at the delamination (yellow symbols) and migration (blue symbols) stages in CB-839 (50 µM) or vehicle (No drug). In A, OCR and ECAR were recorded in individual explants before and after inhibitor treatment and in B-D, mRNAs and ATP levels were measured after two additional hours in the presence of the inhibitor. (E) Scatter dot plots of the intracellular concentrations (mean ± s.e.m.) of glutamine and glutamate in NT explants at the delamination and migration stages. (F) Scatter dot plots of intracellular glutamate concentration (mean ± s.e.m.) in explants at the delamination and migration stages, in CB-839 (50 µM) or vehicle (No drug). (G) Scatter dot plots of the total number of NCCs (left) and of the percentage of Snail-2-positive NCCs (right) per explant (mean ± s.e.m.) after 5 h of NT explants cultured in medium containing either glucose and glutamine or in glucose and glutamate. (H) Scatter dot plot of normalized mRNAs levels (mean ± s.e.m) measured by qRT-PCR of NCC delamination and migration markers in NT explants with quiescent (left) and OXPHOS (right) profiles, in the presence of glutamine or glutamate in the culture medium. (I) Whole mount views of embryos 24 h after injection of glutamate (10 µM) with control siRNAs or siRNAs (28 ng/µl) to GLS and hybridized with probes for *Sox-10*. Detailed views of the trunk region are shown (red bar = 500 µm). The vertical bar delineates the axial levels where NCC development is defective. Arrowheads and arrows point at delaminating and migrating NCCs, respectively. n = 8 for injection of glutamate with control siRNAs with 100% of non-affected embryos and n = 8 for injection of siRNAs to GLS with 75% of the embryos showing apparent defects in NCC delamination and migration. nt, neural tube; so, somite. (J-L) Metabolic parameters (mean ± s.e.m.) of NT explants cultured in medium containing glucose and glutamate including OCR and ECAR (J), the OCR/ECAR ratio (K) and ATP levels (L). (M) Scatter dot plot of normalized mRNAs levels (mean ± s.e.m) measured by qRT-PCR of enzymes of the glutamine pathway in NT explants with a quiescent (left) and an OXPHOS (right) profile, in the presence of glutamine or glutamate in the culture medium. Images in I are from representative experiments. In A, C, I and M, (n) indicate the number of experiments with measurements done in triplicate for each gene and in B, D-G, and J-L, (n) indicate the number of explants analyzed. Data were analyzed by two-way ANOVA relative to the condition with no drug or with glutamine. ns, not significantly different, P > 0.05.

Because GLS converts glutamine to glutamate, we measured the intracellular concentration of glutamine and glutamate in NT explants at delamination and migration after metabolic profiling. Strikingly, the concentrations of glutamine and glutamate were lower at delamination than at migration (Fig. 5E,F). Moreover, the concentration of glutamate in NCCs was severely decreased by CB-839 during migration but not during delamination (Fig. 5E,F). These data suggest that GLS enzymatic activity is low during delamination, but is enhanced during migration, matching the high levels of ATP. To address whether glutamate derived from GLS activity could affect the responses of NT explants, we performed experiments adding glutamate to the culture medium in place of glutamine to bypass GLS enzymatic activity. We found that the number of NCCs, the proportion of Snail-2-expressing cells, and the levels of *Snail-2*, *Foxd-3* and *Sox-10* mRNAs in the presence of glutamate were lower than those measured in the presence of glutamine, indicating that glutamate cannot compensate for the negative impact of glutamine deprivation (Fig. 5G,H; compare with Fig. 3G-I). Likewise, *in ovo* injection of glutamate together with siRNAs to *GLS* in embryos failed to restore the normal NCC delamination and migration patterns (Fig. 5I). Notably, glutamate restored NCC metabolic properties. Indeed, while only an OXPHOS profile was observed in absence of glutamine (Fig. S2A,B), in the presence of glutamate, the NT explants exhibited the two classical OXPHOS and quiescent profiles with characteristics similar to those observed in the presence of glutamine (Fig. 5J-L; compare with Fig. 1B-F). In addition, in glutamate-containing medium the expression of enzymes of glutamine metabolism was altered, with a dowregulation of *GLS* and *GLUL*, and an increase in *GDH*, indicating that glutamate could activate downstream events fueling the TCA cycle (Fig. 5M).

Altogether, our findings reveal that GLS contributes to the NCC delamination process independently from its enzymatic activity, thereby raising the intriguing possibility that GLS exhibits a non-canonical function. In contrast, at the migration stage, GLS enzymatic activity appears to be important for maintaining ATP in association with active locomotion.

### Glutaminase translocates to the nucleus in delaminating NCCs

To identify a potential non-canonical function of GLS during delamination, we compared its cellular localization in delaminating and migrating NCCs. Intriguingly, we found that GLS was particularly enriched in the nuclei of delaminating NCCs in addition to its normal cytoplasmic levels, whereas in migrating cells GLS distributed throughout the cytoplasm without specific nuclear accumulation (Fig. 6A). Treatment of NT explants with CB-839 significantly decreased the proportion of delaminating NCCs exhibiting nuclear GLS but had no effect on GLS localization in migrating cells (Fig. 6A,B). In addition, a similar outcome was observed under glutamine-free conditions in the presence or absence of glutamate (Fig. 6B), suggesting an essential role for glutamine in mediating the nuclear translocation of GLS in delaminating NCCs.

**Figure 6:**
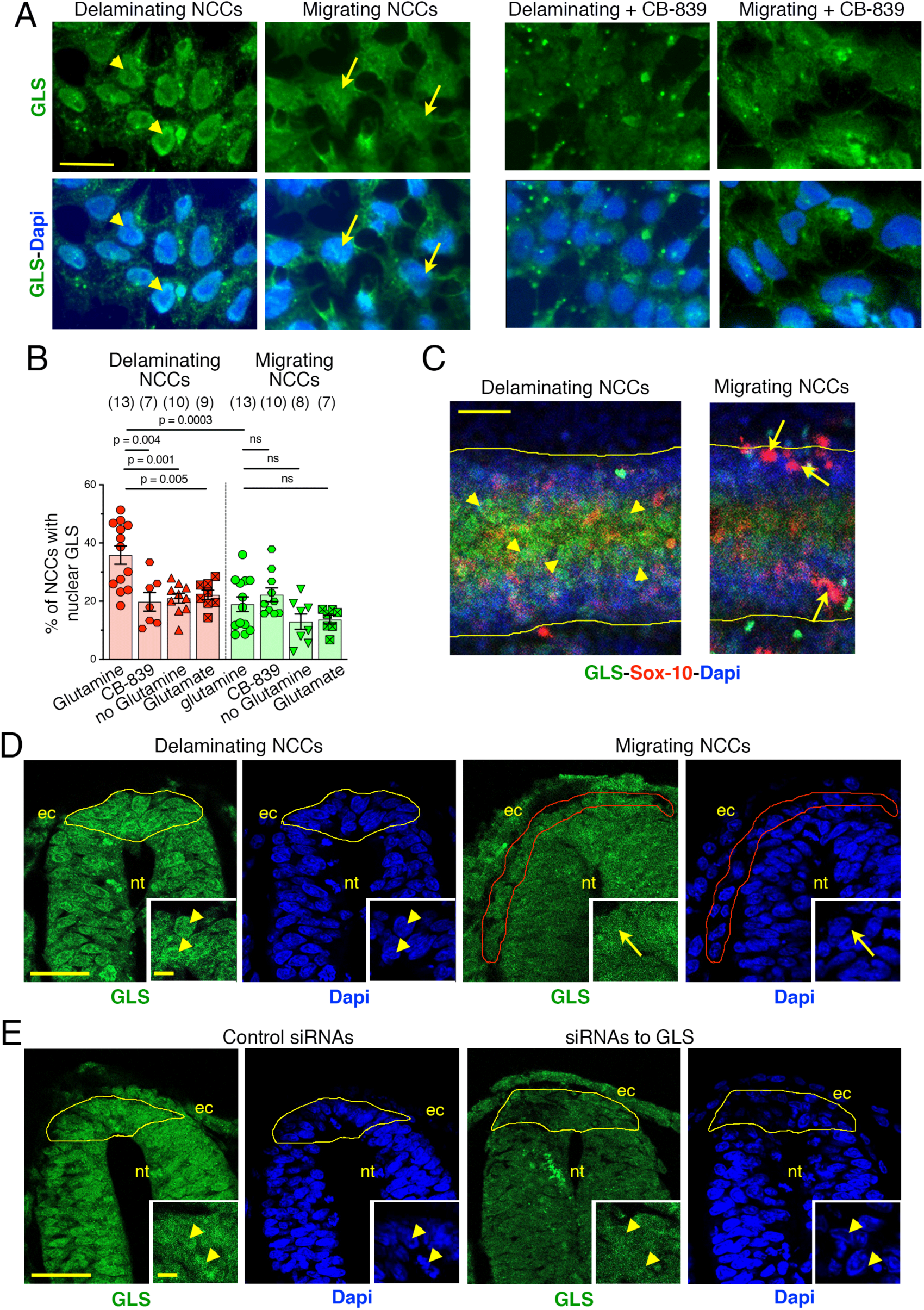
Glutaminase translocates to the nucleus in delaminating NCCs. (A) Immunofluorescence staining for GLS (green) with Dapi visualization of nuclei (blue) in delaminating and migrating NCCs after 5 h in culture in control medium with glucose and glutamine (left) and in medium containing 50 µM CB-839 (right). Yellow arrowheads point at GLS accumulation in nuclei of delaminating NCCs and yellow arrows point at nuclei in migrating cells. Bar = 20 µm. (B) Scatter dot plots of the percentage of NCCs with nuclear GLS (mean ± s.e.m.) among the delaminating and migrating populations in the indicated experimental conditions. CB-839 was applied at 50 µM and glutamate at 10 µM. (C) Horizontal confocal z-projection view of the dorsal part of the NT in the brachial region of a whole mount embryo at the time of NCC delamination (left panel) and early migration (right panel), immunolabeled for GLS (green) and Sox-10 (red) and stained with Dapi (blue); anterior is to the right. Arrowheads point at delaminating NCCs situated along the NT midline and showing nuclear staining for GLS and arrows point at migrating NCCs situated more laterally and showing nuclear staining for Sox-10 but not for GLS. The lateral margins of the NT are delineated with yellow lines. Bar = 50 µm. (n = 3). (D) Confocal sections through the NT in cross-sections at the level of NCC delamination (delineated with a yellow line; left) and of NCC migration (delineated with a red line; right) immunolabeled for GLS (green) and stained with Dapi (blue). Insets show detail of delaminating (left) or migrating (right) NCCs in the dorsal NT. Arrowheads point at the nuclei of delaminating NCCs and arrows point at the nuclei of migrating NCCs. ec, ectoderm; nt, neural tube. Bar = 50 µm and 10 µm in insets. (n = 3). (E) Confocal sections through the NT in cross sections at the level of NCC delamination immunolabeled for GLS (green) and stained with Dapi (blue): left panels, embryos treated for 5 h with control siRNAs at 28 ng/µl; right panels, embryos treated for 5 h with siRNAs (28 ng/µl) against GLS. Insets show detail of delaminating NCCs in the dorsal NT. Arrowheads point at the nuclei of delaminating NCCs. ec, ectoderm; nt, neural tube. Bar = 50 µm and 10 µm in insets. ec, ectoderm; nt, neural tube. Bar = 50 µm. Images in A and C-E are from representative experiments. In B, (n) indicate the number of explants analyzed. Scatter plots were analyzed using one-way ANOVA followed by Dunnett’s multiple comparison tests relative to the control condition in medium with glucose and glutamine without drug. ns, not significantly different, P>0.05.

We also investigated the cellular localization of GLS in trunk NCCs in embryos at delamination stage. Confocal whole mount views of the NT dorsal region revealed the presence of GLS staining within the nuclei of cells situated along the NT midline in the region of NCC delamination, while migratory cells expressing Sox-10 and situated more laterally displayed low GLS nuclear staining (Fig. 6C). Analyses of cross-sections through the NT confirmed the nuclear accumulation of GLS in NCCs at delamination but not at migration (Fig. 6D). Interestingly, in areas where delamination occurs, nuclear localization of GLS was not restricted to NCCs but was also detectable in more ventral NT cells. Finally, we analyzed GLS localization in NCCs of embryos injected with siRNAs to silence *GLS* before delamination. In addition to repressing NCC delamination and expression of EMT-TFs (Fig. 4E-G), *GLS* ablation caused the near disappearance of GLS staining in the cytoplasm and nuclei of NCCs, contrary to control siRNAs, which did not affect GLS expression and intracellular localization (Fig. 6E). Together, our data establish a strong spatiotemporal and functional correlation between GLS expression in the nucleus and its role in controlling delamination of NCCs.

### Glutaminase cooperates with Wnt signaling to control NCC delamination

Delamination of NCCs in the trunk region is under the control of canonical Wnt-1 signaling [38]. The most distinctive step of this pathway is the nuclear translocation of β-catenin, a modular protein either associated with cadherins in intercellular contacts or capable of binding transcription factors in the nucleus to regulate gene expression [39]. Previous investigations in delaminating and migratory NCCs revealed a pattern of nuclear β-catenin localization similar to that observed for GLS in the present study [40], prompting us to explore a possible link between Wnt signaling and a non-canonical function of GLS. We therefore investigated whether β-catenin and GLS are co-distributed in NCCs in culture during delamination and migration. We found that β-catenin was primarily accumulated in the nuclei with GLS in delaminating NCCs (Fig. 7A,B). In contrast, β-catenin was mostly found in cell-cell contacts and rarely in the nuclei of migratory NCCs, while GLS was essentially cytoplasmic (Fig. 7A,B). In embryos, we confirmed GLS localization in the nuclei of NCCs at delamination, while β-catenin showed a diffuse staining in both the cytoplasm and nucleus of cells and was faint in the areas of cell-cell contacts, indicative of a nuclear shuttling activity (Fig. 7C). Importantly, depletion of nuclear GLS by CB-839 *in vitro* or by siRNA *in vivo* caused β-catenin disappearance from the nucleus and cytoplasm of NCCs and its redistribution into cell-cell contacts (Fig. 7B,C), raising the possibility that β-catenin and GLS may be translocated to the nucleus in a coordinated manner. To address this question, we utilized the proximity ligation assay (PLA), to investigate the interaction frequency between β-catenin and GLS. We found a significantly greater number of puncta in the nuclei of delaminating NCCs than in migratory ones (Fig. 7D,E), and this number was markedly reduced in the presence of CB-839, strongly suggesting that nuclear translocation of β-catenin in delaminating NCCs relies on GLS activity (Fig. 7E).

**Figure 7:**
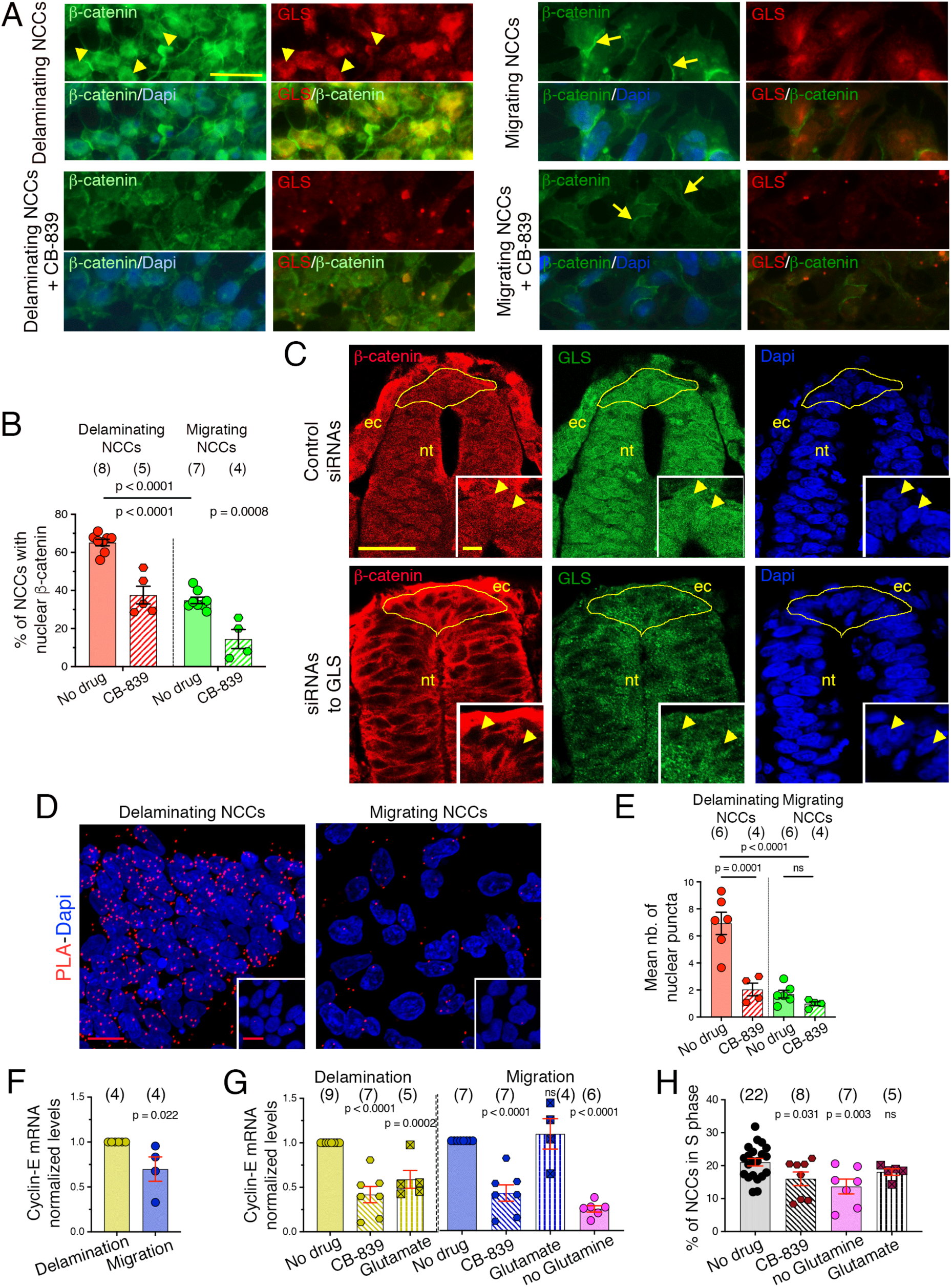
Glutaminase cooperates with Wnt signaling to control NCC delamination. (A) Immunofluorescence staining for β-catenin (green) and GLS (red) with Dapi visualization of nuclei (blue) in delaminating (left) and migrating (right) NCCs after 5 h in culture in control medium containing glucose and glutamine (top) and in the presence of 50 µM CB-839 (bottom). Arrowheads point at β-catenin and GLS accumulation in nuclei of delaminating NCCs and arrows point at β-catenin accumulation in cell-cell contacts of migrating NCCs. Bar = 20 µm. (B) Scatter dot plots of the percentage of NCCs with nuclear β-catenin (mean ± s.e.m.) among the delaminating and migrating populations in the presence or absence of 50 µM CB-839. (C) Confocal images of cross sections through the NT at the level of NCC delamination immunolabeled for β-catenin (red) and GLS (green) and stained with Dapi (blue): top panels, embryos treated for 5 h with control siRNAs (n = 3); bottom panels, embryos treated for 5 h with siRNAs against GLS (n = 3). Insets show detail of delaminating NCCs in the dorsal NT. Arrowheads point at the nuclei of delaminating NCCs. β-catenin pattern is diffuse in the NCC cytoplasm and nuclei coincident with GLS nuclear accumulation in delaminating cells and it is concentrated in cell-cell contacts and absent from the nuclei after siRNAs to GLS treatment. ec, ectoderm; nt, neural tube. Bar = 50 µm and 10 µm in insets. (D) Confocal z-projection views of delaminating (left) and migrating (right) NCCs cultured for 5 h and immunolabeled for β-catenin and GLS followed by PLA treatment (red puncta) and staining with Dapi (blue). Insets show control PLA treatment with GLS antibodies and a non-immune mouse antibody. Bars = 10 µm. (E) Scatter dot plots of the number of nuclear PLA puncta per NCC (mean ± s.e.m.) among the delaminating and migrating populations in the presence or absence of CB-839. (F, G) Scatter dot plot of mRNAs (normalized levels ± s.e.m.) measured by qRT-PCR of Cyclin-E in NT explants at the delamination (yellow) and migration (blue) stages (F) and upon different experimental conditions (G). (H) Scatter dot plot of the percentage of NCCs (both delaminating and migrating) in the S phase (mean ± s.e.m.) in NT explants cultured in the indicated experimental conditions. Images in A, C, and D are from representative experiments. In B, E, and H, (n) indicate the number of explants analyzed, and in F and G, (n) indicate the number of experiments with the measurements done in triplicates for each gene. In B, E, and H, data were analyzed using one-way ANOVA followed by Dunnett’s multiple comparison test relative to the no drug condition. In F and G, data were analyzed by two-way ANOVA relative to the quiescent profile (F) and to no drug condition (G). ns, not significantly different, P>0.05.

In addition to regulating expression of EMT-TFs, Wnt signaling in NCCs drives G1/S transition during the cell cycle, at which step they segregate from the NT [34, 38]. We therefore investigated expression of *cyclin-E*, a member of the cyclin family recruited at the G1/S transition [41, 42], in delaminating and migrating NCCs. We found a higher expression of *cyclin-E* mRNAs in NCCs at delamination than at migration (Fig. 7F). Interestingly, CB-839 negatively affected *cyclin-*E levels at both stages while glutamate decreased *cyclin-E* expression only at delamination (Fig. 7G). Moreover, glutamine deprivation also strongly decreased *cyclin*-*E* expression (Fig. 7G). In agreement with these findings, EdU short-pulse experiments revealed that both CB-839 and glutamine starvation significantly inhibited G1/S transition in NCCs, while glutamate had no effect (Fig. 7H). Altogether, our data indicate that nuclear GLS associates with β-catenin, thus cooperating with Wnt signaling to regulate expression of EMT-TFs and synchronize NCC delamination with the G1/S transition phase of the cell cycle.

## DISCUSSION

Here, we report that NCCs engage different metabolic pathways during delamination and migration in relation with specific gene networks, highlighting that glucose and glutamine are both required for these events, each playing a specific role at each step (Fig. 8). In particular, we show *in vitro* and *in vivo* that during delamination, NCCs exhibit a quiescent metabolic phenotype characterized by a limited energy production due to combined low glycolytic and low mitochondrial respiration activities, yet they are metabolically active. Indeed, delaminating NCCs show a preference for anabolic activity via the PP pathway to support the many molecular changes occurring during EMT. During migration, in contrast, NCCs exploit glucose and glutamine oxidation to fuel mitochondrial respiration for efficient energy production. Thus, NCCs adapt their nutrient preference and use to their bioenergetics and biosynthetic demands to their developmental program. Importantly, this metabolic switch is accompanied by changes in the repertoire of the genes for metabolic enzymes deployed, which is characteristic of a metabolic reprograming. Moreover, we found that metabolic activity not only accommodates to NCC development but also influences their developmental program, as revealed by functional experiments in which blocking glutamine and glucose metabolism affected expression of EMT-TFs and of migration-related genes, causing severe reduction in EMT and migration. Our data therefore uncover a poorly investigated facet of EMT that acquisition of migratory abilities following EMT requires metabolic rewiring in connection with profound cellular and molecular changes. How metabolic pathways are regulated to enable transition from EMT to migration remains yet to be investigated.

**Figure 8:**
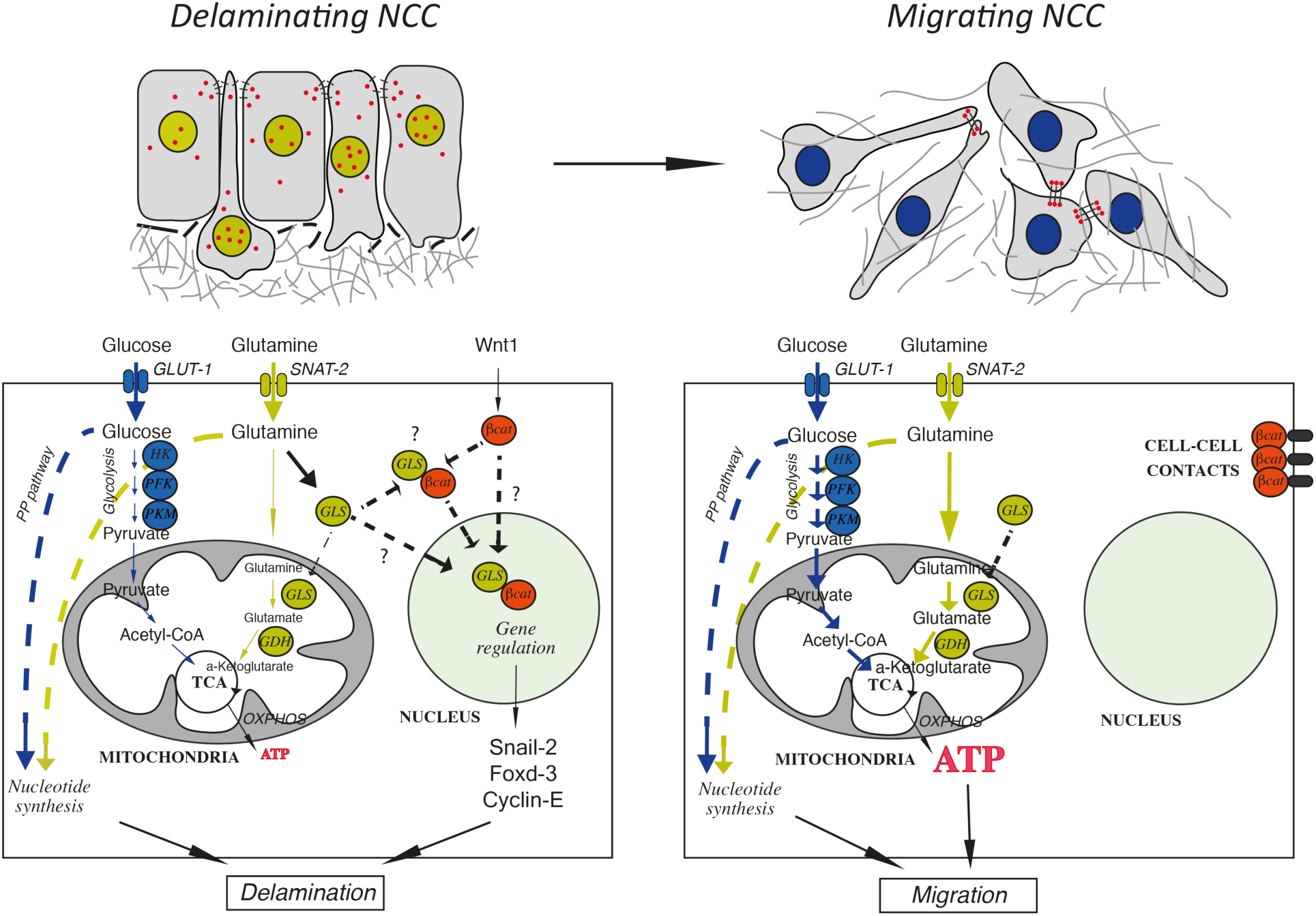
NCC cellular metabolism is coupled to developmental program. Model depicting the roles of the glucose and glutamine metabolic pathways in the control of NCC delamination and migration. Metabolic activity evolves during delamination and migration, from quiescence with reduced production of energy to active mitochondrial respiration for efficient energy production. This shift results in part from a change in glucose utilization from anabolism to active bioenergetics and, more importantly, in the function of GLS, a key enzyme of glutamine metabolism. During delamination, GLS is preferentially translocated into the nucleus under the control of glutamine where it associates with β-catenin (red dots in top panels), a key player of the Wnt signaling pathway. Alternatively, GLS and β-catenin associate in the cytoplasm and are translocated together into the nuclei. Nuclear GLS and β-catenin would participate in the control of expression of key genes, such as *Snail-2*, *Foxd-3* and *Cyclin-E*, to regulate EMT and cell-cycle progression, two necessary events of the delamination process. Once delamination is complete, GLS is no longer present in the nucleus and instead it displays its classical enzymatic activity in mitochondria to convert glutamine into glutamate, to contribute to ATP production together with glucose to promote NCC migration. Coincidently, β-catenin is excluded from the nucleus and is preferentially targeted to cell-cell junctions.

An original aspect and great strength of our work is that we initially identified the metabolic profile of NT cultures prior to performing any analysis of metabolite contribution and gene expression of delamination and migration events. Thus, we could clearly assign a quiescent profile to delamination and an OXPHOS-driven metabolism to migrating cells and confidently manipulate our experimental conditions to investigate the metabolic and molecular mechanisms underlying these two main events in NCC development. The reason for these two distinct metabolic phenotypes in primary cultures that were carefully and identically prepared from embryos handled in the same way is unknown, although we speculate that the *in ovo* environment somehow influences the behavior of cells after isolation.

A most important finding emerging from our study is the signaling role of glutamine during delamination. This function is not mediated directly by glutamine itself but is executed by GLS, a key enzyme of the glutamine pathway. There is increasing evidence that GLS is involved in EMT in cancers [36]. For example, in colorectal cancers, GLS overexpression correlates with increased invasiveness and patient mortality [43]. GLS also has critical role in development of prostate cancers and loss of GLS results in cell cycle arrest and apoptosis [44]. However, the mechanism by which GLS contributes to cell invasion remains elusive and has been mostly attributed to its bioenergetic role. Our data in NCCs advance the understanding of this process, showing that GLS function in EMT is mainly dependent on its nuclear localization and consequent modulation of EMT-TFs expression, while its enzymatic activity leading to glutamate production is of minor importance during this event. This raises the intriguing possibility that GLS exhibits a non-canonical gene regulatory function during NCC EMT. Several examples of nuclear translocation of metabolic enzymes, such as pyruvate kinase M2, an enzyme of the glycolytic pathway [17], have been reported in cancer cells in relation with EMT, but so far this has not been reported for GLS.

A nuclear role for GLS is corroborated by our results demonstrating an interaction between GLS and β-catenin and their concomitant nuclear accumulation in delaminating NCCs. Furthermore, we found that blocking GLS with either an inhibitor (CB-839) or by siRNA abolished the nuclear shuttling of both GLS and β-catenin. Our experiments do not allow us to determine whether GLS and β-catenin associate before nuclear translocation or move to the nucleus separately and interact once there. However, it is worth-mentioning that unlike GLS, β-catenin lacks a nuclear localization sequence in its sequence [45, 46], raising the hypothesis that GLS might serve as a carrier for β-catenin. These findings together with the observation that cyclin-E-dependent G1/S transition is dependent on GLS activity emphasizes a novel critical role of GLS as a master gene regulator in NCC development where canonical Wnt signaling controls multiple events, from induction, EMT, to lineage differentiation. Interestingly, GLS inhibition in prostate cancer has been found to suppress Wnt/β-catenin signaling and inhibit proliferation via repression of cyclin-D1 [44].

The recent demonstration that avian cranial NCCs display a Warburg effect during delamination and that this is coupled with genetic programs controlling cell identity and behavior highlighted the parallel between NCCs and metastasis [3, 23]. However, this view was challenged by the fact that trunk NCCs do not share this feature with cranial NCCs but instead elect a different metabolic process based on glucose oxidation combined with mobilization of multiple metabolic pathways during migration [24]. Our present observation that trunk NCCs utilize nuclear shuttling of GLS as an important intermediate to regulate the gene regulatory network involved in EMT is reminiscent once more of the situation observed in cancer cells. This suggests that despite presenting distinct metabolic features, trunk NCCs share with cancer cells similar processes to connect metabolic events with gene regulation during EMT, establishing that the parallel made between cranial NCCs and cancer cells also applies to trunk NCCs. Finally, these findings illustrates the great diversity of processes used by cells to accomplish their developmental program and raise the possibility that metabolic activity might determine the scenario by which cells undergo EMT.

## MATERIALS AND METHODS

### Reagents

Bovine plasma fibronectin, 1mg/ml stock, was from Sigma (cat. no. F1141). Dispase II at 5 U/ml stock solution in Hanks’ balanced saline was from Stemcell Technologies (cat. no. 07913). The following chemical compounds were prepared and used according to the manufacturers’ guidelines: 2-deoxy-D-glucose (2-DG, Sigma, cat. no. D8375); oligomycin-A (Sigma, cat. no. 75351); rotenone (Sigma, cat. no. R8875); antimycin-A (Sigma, cat. no. A8674); carbonyl cyanide-4-(trifluoromethoxy)phenylhydrazone (FCCP, Sigma, cat. no. C2920); 6-aminonicotinamide (6-AN, Cayman Chemicals, cat. no. 10009315); 6-diazo-5-oxo-L-norleucine (DON, Sigma, cat no. D2141); and CB-839 (Sigma, cat. no. 5337170001). All chemicals were used *in vivo* and *in vitro* at concentrations below those provoking long term-cellular toxicity or overt morphological malformations. 2-DG was solubilized as a 1 M stock solution in DMEM medium (without glucose, pyruvate and glutamine, Gibco, cat. no. A14430-01) and used at 10 mM; oligomycin, rotenone, antimycin, and FCCP were prepared in DMSO as 10 mM stock solutions, diluted as stock solution in culture medium at 100 µM, and used at 1 µM or 0.75 µM (for FCCP); 6-AN was prepared as a 100 mM solution in DMSO and used at 250-500 µM; DON was prepared as a 30 mM stock solution in H_2_0 and used at 30-60 µM; and CB-839 was prepared as a 34 mM stock solution in DMSO and used at 20-50 µM. As 1/1000 dilution of DMSO had not effect on NCC behavior, in experiments using this dilution, the corresponding controls were performed without addition of DMSO. When DMSO dilution was below 1/1000, the same dilution of DMSO but without the drug was used for controls. Four siRNAs corresponding to different regions of the Gallus gallus *GLS* mRNA were designed using the sidirect2.rnai.jp website and purchased from Eurogentec: siRNA.1 GLS: 5’-GAAAGGUUGCUGACUAUAUUC-3’; siRNA.2 GLS: 5’-CCAAAGUUCCUUUUUGUCUUC-3’; siRNA.3 GLS: 5’-GGAUUAAGAUUCAACAAAUUG-3’; siRNA.4 GLS: 5’-AUUUUUCUGCAUUAUUUGCUC-3’. The universal siRNA negative control was purchased from Eurogentec.

### Embryos

Quail embryos were used throughout the study. Fertilized eggs purchased from a local farm (La Caille de Chanteloup, Corps Nuds, France) were incubated at 37-38°C until embryos reached the desired developmental stages. Embryos were staged using the Hamburger and Hamilton (HH) chart [33] and based on the number of somite pairs.

### *In ovo* injection of drugs and siRNAs

I*n vivo* loss-of-function experiments were performed on embryos at stage HH12 (15-16 somite pairs at 48 h of incubation), i.e. before onset of delamination. After removing 1 ml of albumin from the egg and opening the shell with curved dissecting scissors to expose the embryo to the experimenter, a drop of Fast Green solution was added on top of the vitelline membrane for easy visualization of the embryonic tissues, and the drugs and siRNAs diluted at the desired concentration in PBS were injected using glass pipets at the tail bud end of the embryo into the lumen of the NT and under the vitelline membrane in the vicinity of the NT over the unsegmented region and the last 3-5 somites (10 μl delivered per injection). Controls embryos were injected with the vehicle only. The eggs were sealed with tape and were further incubated at 37-38°C in a moist atmosphere for 5-24 h. Monitoring of NCC development was performed by following expression of marker genes by qRT-PCR at 5 h and whole mount in situ hybridization at 5 and 24 h followed by sectioning (see below).

### Cell culture

#### Generation of NCC primary cultures

NCC cultures were produced from trunk NT obtained from quail embryos at different stages covering the delamination process: stage HH12-13 (17-18 somite pairs), stage 13 (19-21 somite pairs) and HH14 (23-25-somite stage) as described [24, 35]. After opening the eggshell, the yolk was transferred into phosphate-buffered saline (PBS). The embryo was cut off from the yolk and transferred into PBS in an elastomer-containing dish. An embryo portion of about 750-µm long was excised at the level of the last 5 somites with a scalpel under a stereomicroscope and subjected to mild enzymatic digestion by treatment with dispase II at 2.5 U/ml for 5-10 minutes at room temperature. The NT was dissected out manually using fine dissection pins under a stereomicroscope, freed from the surrounding tissues, and transferred for 30-60 minutes in DMEM medium without glucose, pyruvate and glutamine (Gibco, cat. no. A14430-01) supplemented with 0.5% fetal bovine serum, for recovery from enzyme treatment. NTs were explanted onto a variety of substrates depending on the aim of the culture (plastic culture dishes, Seahorse plates, chambered glass coverslips or glass coverslips), all coated with fibronectin at 10 µg/ml in PBS (i.e. about 5 µg/cm^2^) for a minimum of 1 h at 37°C. To ensure rapid initiation of NCC migration, NTs were positioned with their dorsal side oriented down toward the substratum. Explants were cultured at 37°C under normoxic conditions in a 5%-CO_2_ incubator in DMEM containing 1% serum, 100 U/ml penicillin, 100 µg/ml streptomycin, and supplemented or not with 5 mM glucose, 2 mM glutamine or both, depending on the aim of the experiment. In some experiments, glutamine was replaced by glutamate (Sigma, cat. no. 49621) used at 10 µM. The NT explants were followed and analyzed routinely during the first 5 h of culture and up to 24 h in some experiments. Throughout each experiment, the morphology of the NT explant, area and progression of the NCC outgrowth, as well as individual cell shape were evaluated, imaged, and assessed regularly under an inverted phase contrast Nikon microscope equipped with 6.3X, 10X, and 20X objectives.

#### Quantifications of NCC numbers

To quantify the number of NCCs in outgrowths, phase contrast images of explants encompassing the whole NCC outgrowth were taken with the 10x objective at defined time points of the culture. Cells were counted in a square of 100-µm-side of the outgrowth using ImageJ and data were reported to the entire outgrowth area measured using ImageJ. To analyze the delaminating and migrating NCC populations in NT explants in culture at the end of the experiments, the NT was removed manually from the dish by gentle flushing of culture medium with a pipet tip, thereby uncovering delaminated cells situated underneath the NT [24]. Delaminating cells were discriminated from migratory cells using several criteria: their strong Snail-2 content, their central position in the outgrowth often separated from the migration zone by a cell-free gap, their reduced size, and their compact shape. The numbers of delaminated and migrating cells were counted as above in a square of 100-µm side covering the areas of delaminated cells or of migrating cells, in the mid-part of the explant along its long axis and reported to the entire outgrowth area.

### Cellular bioenergetics analyses

#### Seahorse analyses

Bioenergetics profiles of NCC primary cultures were determined using a Seahorse Bioscience XF24 Analyzer as described [24]. A single NT explanted from embryos at HH13 was deposited precisely at the center of each well of a 24-well Seahorse plate previously coated with fibronectin and containing 100 µl of culture medium, and NCCs were allowed to undergo migration for 2 h (Fig. S1B). Before preparation of the plate for Seahorse analysis, phase contrast pictures of the NT explants were taken and their areas were measured for normalization. Cultures were then rinsed in Seahorse XF medium (Agilent, cat. no 103575-100) without serum and incubated for 30 minutes at 37°C in normal atmosphere in 500 µl assay medium supplemented with the same metabolites as prior to the rinsing step, except serum. Then, the Seahorse assay was run according to the manufacturer’s instructions. After Seahorse metabolic profiling, NCC primary cultures were processed for cellular assays of delamination and migration, immunolabeling, qPCR analyses, or intracellular ATP, glutamine or glutamate measurements (Fig. S1B). All experiments were normalized with respect to both the NT length and the stage of embryos at onset of experiments.

Oxygen consumption rate (OCR) and extracellular acidification rate (ECAR) values, as readouts of basal mitochondrial respiration and lactate production, were assessed through 4 cycles of measurement during 30 minutes, and in some experiments, this was followed by drug injections and 3 additional cycles of measurement. All drugs solutions were prepared from stock solutions in assay medium supplemented with glucose and glutamine. OCR and ECAR values were used to establish the energetic maps of cells, comprising four quadrants (quiescent, aerobic (OXPHOS), glycolytic, and energetic) as described previously [24]. The key parameters of mitochondrial respiration (ATP-linked respiration, maximal respiratory capacity and proton leak) were measured by means of a MitoStress test after sequential additions of oligomycin at 1 µM, FCCP at 0.75 µM, and rotenone + antimycin at 1 µM through 3 cycles of measurements in 30 minutes. Coupling efficiency, which represents the proportion of respiratory activity that is used to make ATP, was calculated as the difference between the basal OCR and the minimal, ATP-linked OCR. Spare respiratory capacity, which corresponds to the extra mitochondrial capacity available in a cell to produce energy under conditions of increased demand, was calculated as the difference between the maximal OCR and the basal OCR [47-49].

Temporal metabolic changes of NTs were investigated using two rounds of Seahorse assays on the same explants. After identifying the initial metabolic profile of NT explants at the first 3 h of culture, the plate was removed from the analyzer, the assay medium was changed for 100 µl of fresh culture medium and the plate was further incubated for overnight at 37°C in a 5%-CO_2_ incubator and then proceeded for the second run of Seahorse assay under the same conditions as for the first one.

#### ATP and metabolite measurements

Intracellular ATP and glutamine/glutamate levels were measured for each NCC primary cultures after Seahorse analysis using the ATPlite Bioluminescence assay kit from PerkinElmer (cat. no. 6016943), and the Glutamine/Glutamate-Go assay (J8021) assay kit from Promega (cat. no. J8021), respectively, following the manufacturers’ instructions. Measures were normalized to NCC numbers.

### Cryosectioning

To obtain embryo sections for immunolabeling or following in situ hybridization, embryos were washed in PBS for 1 h at room temperature and in 15% sucrose solution overnight at 4°C. Next, they were incubated in 7.5% porcine gelatin (dissolved in 15% sucrose solution) for 2-3 h at 40°C, embedded in gelatin-sucrose in cup, snap-frozen in chilled isopentane at -70°C and stored at -80°C. 20-µm sections were obtained using the Leica cryostat and collected on Superfrost/Plus slides (Thermo Fisher Scientific). For imaging, the slides were immersed in PBS at 40°C for 2 h for gelatin removal, washed in PBS and mounted in Aquatex mounting medium (Merck). Sections of embryos with in situ hybridization were observed and imaged with a Hamamatsu NanoZoomer Digital Slide Scanner and treated with NDP.View2 software. Immunolabeled sections were observed with confocal microscope 20X fluorescence objectives.

### Immunofluorescence labeling of NCC cultures and embryo sections

For immunolabeling, the following primary antibodies were used: Rabbit monoclonal antibody (mAb) to Snail-2 (clone C19G7, Cell Signaling, 1/300), mouse mAb to Sox-10 (Clone A2, Santa Cruz, 1/200), mouse mAb to β-catenin (clone 14, BD-Transduction Laboratories, 1/200), and rabbit polyclonal antibodies to GLS (PA535365, Life Technologies, 1/50). NCC primary cultures were performed on glass coverslips deposited in 4-well plates (Nunc, cat. no. 144444) or in 8-well Lab-Tek Chambered Coverglass (Nunc, cat. no. 154511) coated with fibronectin. After removing the NT to uncover delaminating NCC, the cultures were fixed in 4% paraformaldehyde (PFA) in PBS for 15 minutes at room temperature for detection of all antigens, except β-catenin (45 minutes in 1.5% PFA). After permeabilization with 0.5% Triton X-100 in PBS for 5 minutes and washes in PBS, cultures were blocked in PBS-3% BSA and subjected to immunofluorescence labeling using primary antibodies followed by incubation with appropriate secondary antibodies conjugated to Alexa-fluor 488 or Cy-3 (Jackson Immunoresearch Laboratories), and processed for DAPI or Hoechst staining to visualize cells’ nuclei before mounting in ImmuMount medium (Shandon). For immunostaining of embryo sections, essentially the same procedure was used. Preparations were observed with a Zeiss AxioImager M2 epifluorescence microscope equipped with 20X-63X fluorescence objectives (Acroplan 10X/0.25, 20X/0.45, Plan-Neofluar 40X/0.75 and 63X/1.25 oil) or using z-stack acquisitions with a Zeiss LSM 900 confocal microscope equipped with 10X-40X fluorescence objectives (Plan-Apochrome 40X/1.30 oil). Data were collected using the Zen or Airyscan2 software and processed using ImageJ software.

### Whole mount immunolabeling

Trunk regions of embryos were fixed with 4% PFA for 2 hours at 4 C°, washed 2 times with PBS + 0.1% Triton X100 for 5 minutes, and incubated with blocking solution (PBS + 0.1% Triton X100 + 1% BSA and 0.15 % (w/v) glycine) for 2 days at 4 C° under stirring. They were then incubated for one night at 4 C° with gentle shaking with primary antibodies to GLS (1:50) and Sox-10 (1:200) in blocking solution. In the following day, the embryos were washed 3 times for 20 minutes with blocking solution at room temperature and then incubated for another night with secondary antibodies conjugated to Cy3 and Alexa-fluor 488 (1:300) in blocking solution as for the primary antibodies. Lastly, embryos were washed 3 times for 20 minutes with blocking solution and transferred into a Coverwell imaging chamber (Grace Bio Lab, cat. no. 635041) with their dorsal side oriented toward the bottom. After removing the bulk of liquid to keep them flat, a few droplets of mounting medium were added before mounting with a coverslip, and the preparations were kept for 24 hours at room temperature before imaging with confocal microscope 20X fluorescence objectives.

### Proximity ligation assay (PLA)

After 5 hours in culture and removing the NT to uncover delaminating NCC, explants were fixed with 4% PFA for 15 minutes, followed by permeabilization with 0.5% Triton X-100 in PBS for 5 minutes, rinsing in PBS and blocking in PBS-3% BSA. Cultures were then subjected to immunofluorescence labeling using pairs of primary mouse and rabbit antibodies to β-catenin and GLS diluted in blocking solution as described above. Non-immune mouse antibodies (Jackson ImmunoResearch, cat. no 015-000-002) were used instead of mouse mAb to β-catenin for controls. Following primary antibodies incubation, the cultures were washed 3 times in PBS for 5 minutes and incubated with Duolink Plus and Minus PLA probes (diluted 1:5) followed by Duolink In Situ Orange Starter Kit Mouse/Rabbit (Sigma, cat. no. DUO92102), according to the manufacturer’s guidelines. Confocal microscopy image acquisition was performed as above and quantitative analysis of PLA-positive dots was done on Z-projection images through the entire cell using ImageJ.

### In situ hybridizations for mRNA detection on whole mount embryos

In situ hybridization mRNA probes were generated either by PCR (for *GLS* and *GDH*) or from plasmid vectors (for *Sox-10* and *Foxd-3*). For PCR probes, primers were designed to generate a 400-500 bp fragment of the mRNA coding sequence based on information available from NCBI databases. The following forward and reverse sequences were selected for GLS, *fwd* 5’-GAAGGACAAGAGAAAATACCAGTG-3’, *rev* 5’-TGCTCCAGCATTCACCATAGG-3’; GDH, *fwd* 5’-TGGGACAATCATGGGCTTCC-3’, *rev* 5’-CAGTACGCATGATTTGTCGAGC-3’; and SNAT-2, *fwd* 5’-TACGCTGTCCCAATCCTGAC-3’, *rev* 5’-TGCTGCAGATGCACCAATGAAT-3’ and the PCR primers were designed as follows. The linker, 5’-GAG-3’, and the sequence of the RNA polymerase T7 binding site, 5’-TAATACGACTCACTATAGGG-3’, were added to the reverse sequences. The synthetized PCR primers (Eurogentec) were used to generate SNAT-2, GLS, and GDH probes by PCR reaction performed with chick embryo cDNA template using Platinum Taq DNA polymerase (Invitrogen, cat. no. 10966018). PCR products were assessed on a gel and purified using the Qiaquick PCR Cleanup kit (Qiagen, cat. no. 28104). Plasmids for *Foxd-*3 and *Sox10* mRNA probe synthesis were from C. Erickson and P. Scotting, respectively [50, 51]. Linearized plasmid DNA and PCR products were used to synthesize digoxigenin-UTP (Roche) labeled antisense probes with RNA polymerases from Promega and RNA probes were purified with Illustra ProbeQuant G-50 microcolumns (GE Healthcare, cat. no. 28903408).

In situ hybridizations were performed on whole mount embryos collected at the appropriate developmental stages, and fixed in 4% PFA in PBS for 2 h at room temperature or overnight at 4°C. Embryos were hybridized overnight at 65°C with the digoxygenin-UTP-labeled RNA probes in 50% formamide, 10% dextran sulfate and Denhart’s buffer (0.5 µg probe/ml hybridization buffer) and washed twice in 50% formamide, 1x SSC and 0.1% Tween-20 at 65°C, then 4 times at room temperature in 100 mM maleic acid, 150 mM NaCl pH 7.5 and 0.1% Tween-20 (MABT buffer). After a 1-h pre-incubation in MABT buffer containing 10% blocking reagent (Roche) and 10% heat-inactivated lamb serum, embryos were incubated overnight at room temperature with the anti-digoxygenin antibody (Roche). After extensive rinsing with MABT buffer, they were preincubated in 100 mM NaCl, 50 mM MgCl_2_, 1% Tween-20, and 25 mM Tris-HCl, pH 9.5, and stained with NBT-BCIP (Roche) following manufacturer’s guidelines. Preparations were observed and imaged with a Nikon stereomicroscope.

### mRNA quantification by quantitative RT-PCR analyses

For NT explants cultured in vitro, after characterizing the metabolic profiles of individual explants by Seahorse analysis, total RNA of explants sharing the same metabolic profile were isolated using Trizol (Invitrogen), following the manufacturer’s instructions. For normal, untreated embryos and for embryos injected with drugs or siRNAs, the NT situated in the caudal part encompassing the last 5 somites up to the tail bud was dissected manually with fine dissection pins, and the total RNA was extracted using Trizol (Fig. S1C, D). 500 ng RNA were used for cDNA synthesis using SuperScript IV reverse transcriptase (Invitrogen, cat. no. 18090050). Quantitative real-time PCR were performed using the Power Syber Green Master Mix (Applied Biosystems, cat. no. 4368708) in a StepOne Plus RT-PCR apparatus (Applied Biosystems). Gene expression was assessed by the comparative CT (ΔΔ_Ct_) method with β-actin as the reference gene.

The following primers were used:

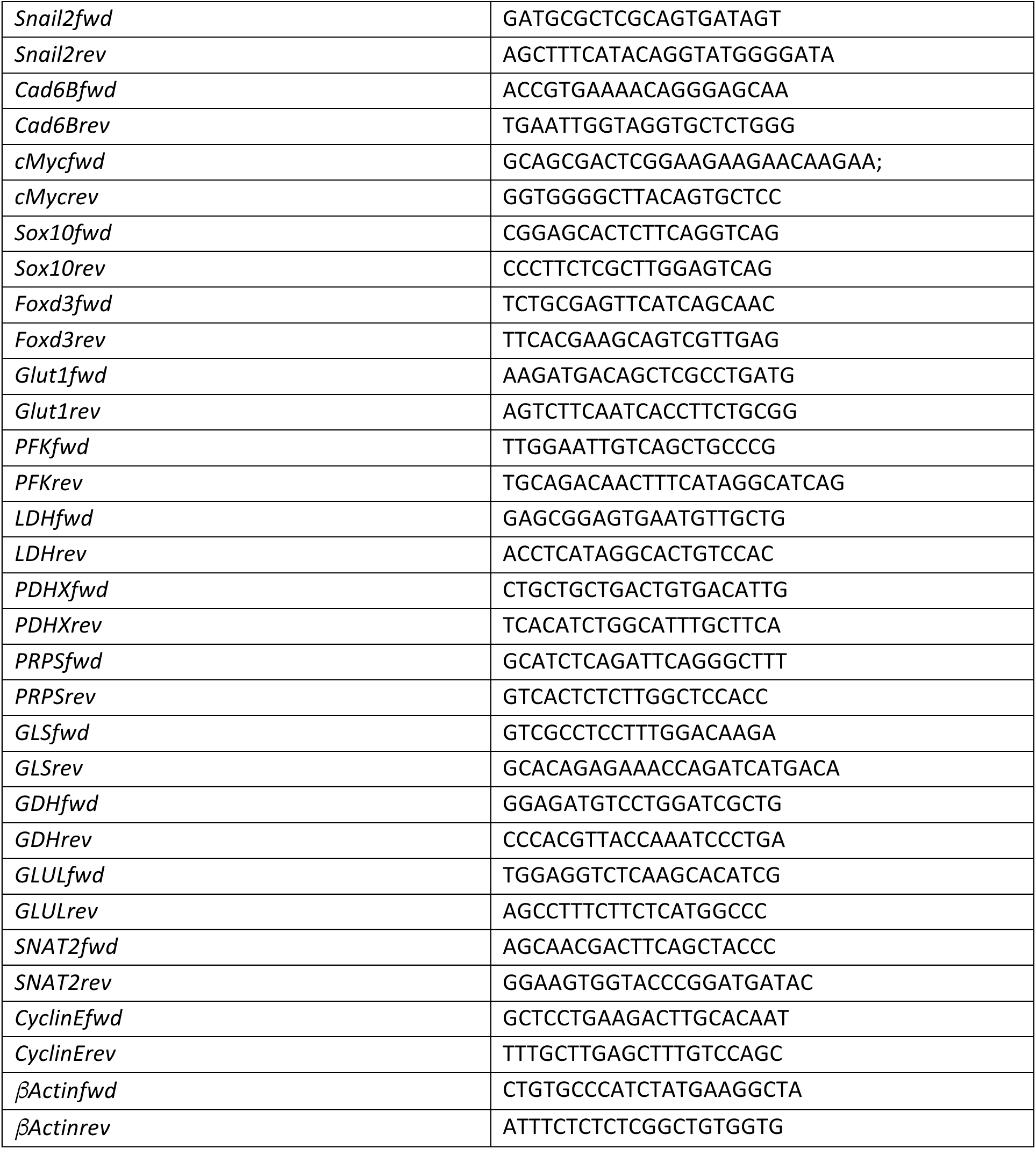

### Cell Cycle Assay

Cell cycling in culture was monitored using the Plus EdU Cell Proliferation Kit for Imaging from Life Technologies (cat. no. C10638). Briefly, NCC primary cultures were incubated with EdU at 20 µM in culture medium for 1 h. Immediately after EdU incorporation, cultures were fixed in 4% PFA in PBS for 15 minutes at room temperature, permeabilized in 0.5 Triton-X100 for 15 minutes and treated for EdU detection using Alexa-555 Fluor azide in accordance with the manufacturer’s guidelines. After DNA staining with Hoechst, cultures were examined as described for immunolabelings.

### Cell locomotion assay

NCC primary cultures were performed in 4-well Chambered Coverglass (Nunc, cat. no. 155383) coated with fibronectin. Up to 5 NT were distributed separately into each well, and were maintained at 37°C in a humidified 5%-CO_2_ incubator for about 2 h until the NT adhered firmly to the dish and NCC initiated migration on the substratum; then the cultures were transferred into a heated chamber (Ibidi) with a humid atmosphere containing 5% CO_2_/95% air placed on the motorized stage of a Leica DMIRE2 microscope equipped with a CoolSNAP HQ camera (Roper Scientific). Time-lapse video microscopy was performed with a 10X objective and phase contrast images were captured every 5 minutes during 16-24 h using the Micromanager software. Trajectories and positions of individual NCC in several NT explants recorded in parallel were tracked using Metamorph 7 software. The progression of the migration front of the NCC population every hour was measured using a custom GUI (available upon request) written in MATLAB (MathWorks., Natick, MA, USA). Trajectories and positions of individual NCCs in several explants recorded in parallel were tracked using Metamorph 7 software. The velocity of each NCC was calculated as the ratio between the total length of its trajectory and the duration of the acquisition time.

### Statistical methods

Statistical analyses were performed using Prism 7 (GraphPad). For statistical analysis of data, we used One-way ANOVA parametric test after the validation of normality and equality of variances using Shapiro-Wilk and Brown-Forsythe methods, respectively. For comparison between two conditions we used unpaired t-test two-tailed test after the validation of normality and equality of using Shapiro-Wilk and equality of variances. Otherwise, non-parametric Mann-Whitney test was used. Unless specified, at least three independent experiments were carried out for each procedure. Each NT explant was considered as an individual sample. The data obtained for each drug or nutrient condition were compared to that of the medium without drug or with both nutrients, respectively. The number of samples n (explants or embryos) analyzed is indicated in each graph in brackets. For statistical analysis of qRT-PCR results of in-vitro experiments, mRNA expression from a pool of 1-8 NT explants are presented as mean ± s.e.m of triplicates per experiment and gene. In *in-vivo* experiments, mRNA expression was measured per embryo after analyzing of three replicates measurements acquired per embryo and genes. The data obtained from embryos injected with drugs were analyzed using Two-way ANOVA relative to the data of embryos injected with vehicle. Data are expressed as mean values ± s.e.m. Results are considered statistically significantly different when p<0.05.

## ACKNOWLEDGMENTS

We deeply thank Chantal Thibert and Sakina Torch for providing advice in cellular metabolism. Special thanks to Redouane Fodil for designing the software to track the progression of the migratory front of NCC populations. Many thanks to Xavier Decrouy from the IMRB imaging platform and Xavier Laffray from the histology and imaging platform of the Laboratoire Gly-CRRET for advice.

## COMPETING INTERESTS

The authors declare no competing financial interests.

## AUTHOR CONTRIBUTIONS

N.N.-M. conceived the project and designed the experimental approach, performed experiments, analyzed data, and contributed to the manuscript. A.D. provided expertise for *in ovo* studies and facilities for cryo-sectioning methods and analyses. F.R. provided financial and logistics support. R.F. and R.M. provided expertise in cellular metabolism and Seahorse technology and revised the manuscript. S.D. designed the experimental approach, performed experiments, analyzed the data, and contributed to the manuscript. J.-L.D. designed the experimental approach, performed experiments, analyzed the data, and wrote the manuscript.

## FUNDING

This work was supported by funding from Institut National de la Santé et de la Recherche Médicale, Université Paris-Est Créteil, AAP IMRB cross-teams project, and Fondation ARC pour la Recherche sur le Cancer (No. PJA 20181207844). N.N.-M. was funded by doctoral fellowship of Université Paris-Est Créteil and by the Labex REVIVE.

## EXTENDED DATA FIGURES

**Extended data Figure 1:**
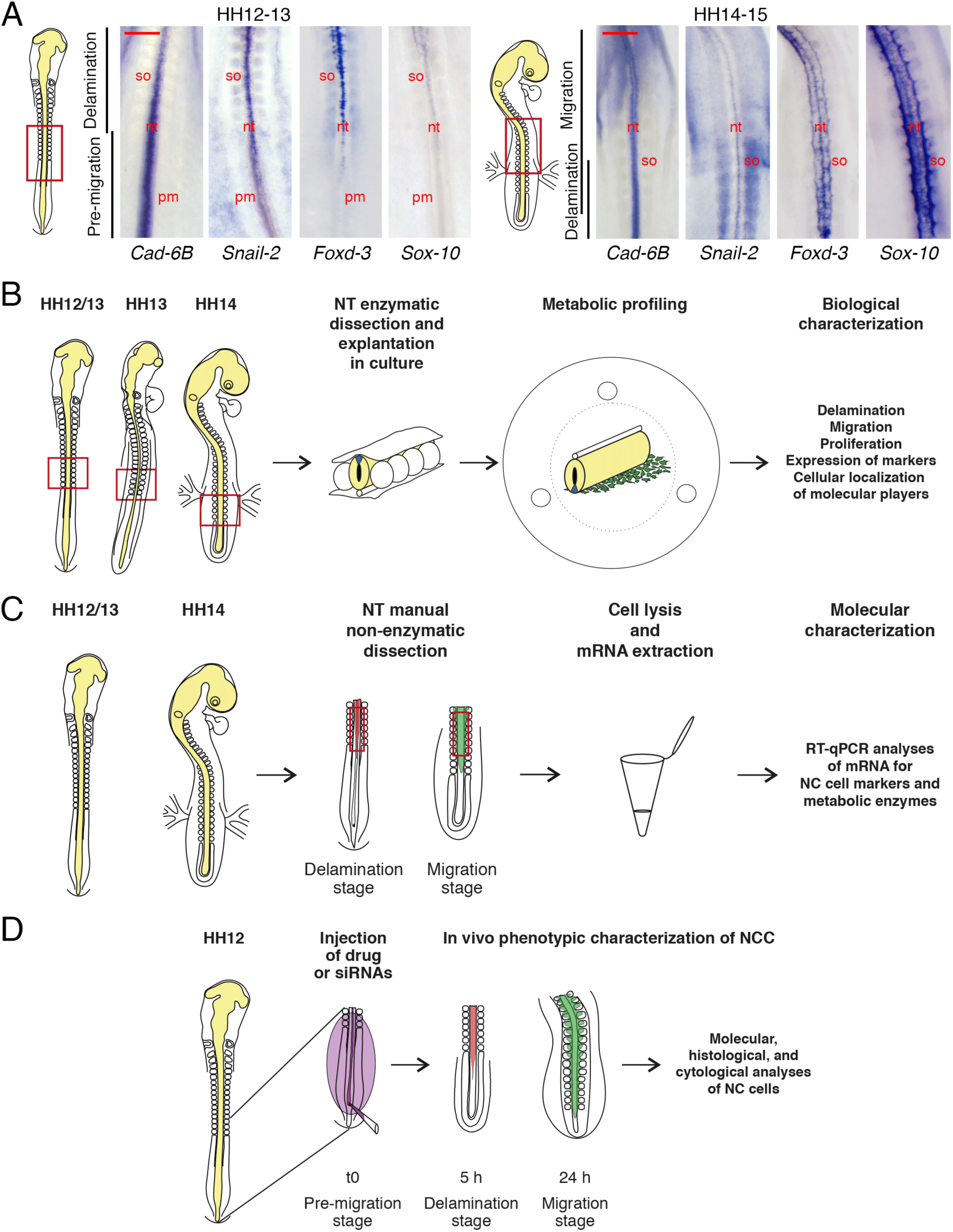
(A) Expression patterns of markers for NCC delamination and migration at stages HH12-13 (left) and HH14-15 (right). Whole mount views of the thoracic region at mid-trunk level (corresponding to somite pairs 15-22) of quail embryos hybridized with probes for *Cadherin-6B*, *Snail-2*, *Foxd-3*, and *Sox-10*. The diagrams depict the gross morphology of the embryos at the respective stages, and the regions shown in the in situ hybridization images are delineated by red rectangles. In situ hybridizations for each probe and at each developmental stage were done in triplicate. Bar = 500 µm. nt, neural tube; pm, presomitic paraxial mesoderm; so, somite. (B) Schematic representation of the procedure for analyzing *in vitro* the metabolic profile of individual NT explants at stages HH12/13 to HH14 and the corresponding molecular and cellular properties of NCCs (see Methods for details of the procedure). The embryonic region extirpated for NT dissection is delineated with a red rectangle. (C) Schematic representation of the procedure to characterize the molecular features of NTs and associated NCCs collected from embryos at stages HH12-13 and HH14. The embryonic region extirpated for mRNA extraction is delineated with a red rectangle. (D) Schematic representation of the procedure for *in-ovo* injection of quail embryos for loss-of-function studies with indication of the timing of NCC development (see Methods for details of the procedure). In C and D, zones with NCC at the delamination and migration stage are colored in red and green, respectively. Images in A are from representative experiments.

**Extended data Figure 2, related to Fig. 3:**
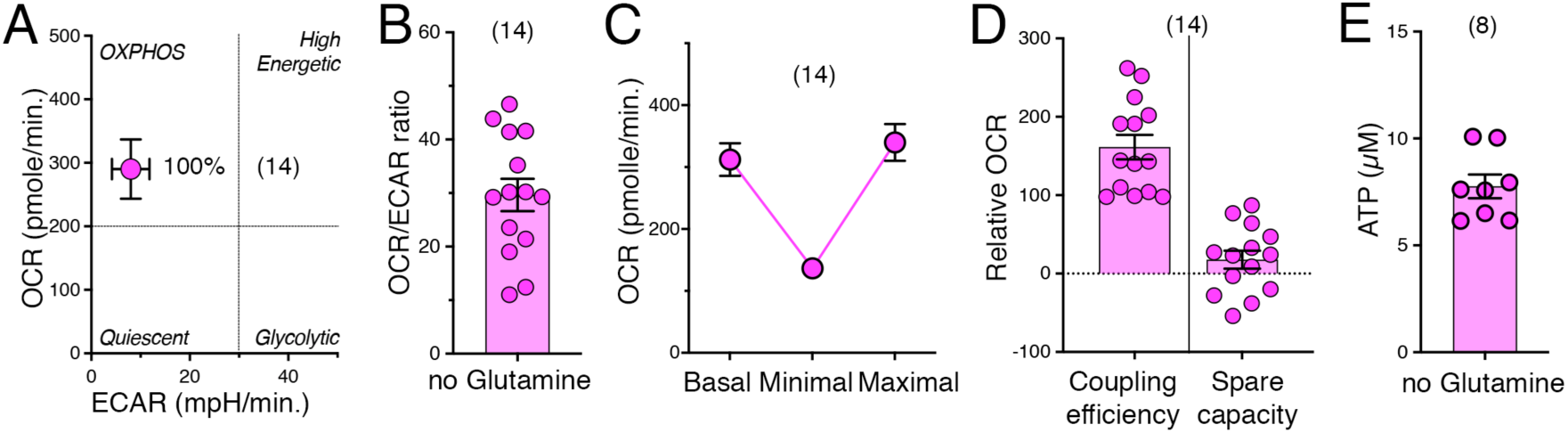
(A-E) NT explants cultured in medium lacking glutamine (magenta symbols) were analysed for metabolic parameters (mean values ± s.e.m.) including OCR and ECAR (A); the OCR/ECAR ratio (B); basal, minimal, and maximal OCR (C); coupling efficiency (left) and spare respiratory capacity (right) (D); and ATP levels (E). n indicates the number of explants analyzed.

## Notes

### Competing Interest Statement

The authors have declared no competing interest.

